# Vectashield-induced fluorescence quenching hinders conventional and super resolution microscopy

**DOI:** 10.1101/393264

**Authors:** Aleksandra Arsić, Nevena Stajković, Rainer Spiegel, Ivana Nikić-Spiegel

## Abstract

Finding the right combination of a fluorescent dye and a mounting medium is crucial for optimal microscopy of fixed samples. It was recently shown that Vectashield, one of the most commonly used mounting media for conventional microscopy, can also be applied to super-resolution direct stochastic optical reconstruction microscopy (dSTORM). dSTORM utilizes conventional dyes and starts with samples in a fluorescent ON state. This helps identifying structures of interests. Subsequently, labelled samples are brought to blinking, which is necessary for localization of single molecules and reconstruction of super-resolution images. This is only possible with certain fluorescent dyes and imaging buffers. One of the most widely used dyes for dSTORM, Alexa Fluor (AF) 647, blinks in Vectashield. However, after adding Vectashield to our samples, we noticed that the fluorescence intensity of AF647 and its improved variant, AF647+, is quenched. Since structures of interest cannot be identified in quenched samples, loss of fluorescence intensity hinders imaging of AF647 in Vectashield. This has consequences for both conventional and dSTORM imaging. To overcome this, we provide: 1) a quantitative analysis of AF647 intensity in different imaging media, 2) practical advice on how to use Vectashield for dSTORM imaging of AF647 and AF647+.

## 1. Introduction

Super-resolution microscopy (SRM) provides unique insights into the subcellular organization of cells and tissues. Several techniques have been developed to make the nanoscale imaging possible^1-4^. Single molecule localization techniques, such as photoactivated localization microscopy (PALM)^5^, fluorescence photoactivation localization microscopy (FPALM)^6^, stochastic optical reconstruction microscopy (STORM)^7^ and direct stochastic optical reconstruction microscopy (dSTORM)^8^ rely on temporal separation of fluorescence emission. Temporal separation is achieved by blinking, which separates individual fluorescent molecules not only in time, but also in space. The resulting spatiotemporal separation of fluorescence emission enables precise localization of fluorescent molecules, which is necessary to reconstruct SRM images^7,9^. While PALM and FPALM utilize genetically engineered fluorescent proteins, dSTORM relies on synthetic fluorophores (dyes) and conventional labelling techniques. To achieve blinking in dSTORM, dye molecules are switched to a dark OFF state from which they stochastically and spontaneously recover to a fluorescent ON state, multiple times before bleaching. Blinking can be induced in certain dyes by using strong lasers and particular imaging buffers, such as thiol-containing reducing agents with or without enzymatic oxygen scavenger systems (e.g. GLOX buffer containing glucose oxidase and catalase)^8,10^.

While blinking is essential for dSTORM, having the dye molecules in the fluorescent ON state before the induction of blinking is of high importance for optimal dSTORM imaging. Being able to see the fluorescent signal is necessary for the identification of positively labelled cells and structures of interest. This is relevant for experiments based on transfections in which depending on the transfection efficiency, only a certain number of cells will be transfected and thus can be imaged under a microscope. This is equally relevant when doing immunocytochemistry or immunohistochemistry experiments, in which specific regions of the cells or tissues need to be identified. In addition, the usual practice in dSTORM and other SRM techniques is to take a reference image with conventional diffraction limited microscopy prior to switching to the SRM mode. For this, fluorescent molecules also need to be in the ON state first. Furthermore, having the dye molecules in the fluorescent ON state helps with judging the quality of labelling since only with molecules in the ON state signal-to-noise ratio can be measured. Last but not least, having molecules in the ON state is necessary for setting a correct focus point for 3D dSTORM imaging^11^.

It was recently reported that Vectashield, a commercial mounting medium for confocal microscopy, can also be used as an imaging buffer for structured illumination microscopy (SIM)^12^, dSTORM^13^ or their combination^14^. In comparison with self-made buffers, commercially available Vectashield represents a very convenient and affordable choice for dSTORM imaging medium. The fact that it can be used off-the-shelf is particularly beneficial for demanding techniques such as dSTORM. Using the same commercially available imaging medium brings more reproducibility and allows for more comparable results within and between different laboratories. In addition, Vectashield has high refractive index and can be used for imaging with objectives with high numerical apertures. Another advantage is that in contrast to GLOX buffer, Vectashield is more stable and can be used for longer imaging sessions. Due to the depletion of oxygen scavenging system, GLOX buffer can only be used for a couple of hours. On the other hand, samples can stay in Vectashield for several days^13^.

The first report on the use of Vectashield for dSTORM studied Vectashield’s suitability for dSTORM imaging^13^. The authors imaged microtubules and tested several fluorescent dyes, such as Alexa Fluor (AF) 647, AF700, AF555, Cy3. The best blinking was achieved with AF647. Combining Vectashield and AF647 for dSTORM has the highest practical importance since AF647 is the most widely used dye for dSTORM. However, this is in contrast to one previous report on the incompatibility of AF647 and Vectashield^15^. While Olivier and colleagues also point out that Vectashield is not suitable for every dye, they do not mention any potential issues when imaging AF647 in Vectashield, nor is it mentioned in other studies that used Vectashield for AF647 imaging^14,16-18^.

The reason for this could be that until now just a few cellular targets were imaged with dSTORM in Vectashield – mainly cytoskeletal elements, such as actin and microtubules. These structures are relatively easy to image and usually serve as proof-of-principle target proteins when comparing different SRM techniques or imaging conditions. In our hands, imaging of these proteins in Vectashield was also straightforward, but this was not the case when using Vectashield for imaging of less abundant proteins. In the latter case, the problem was that we could not identify any of the AF647-labelled targets in Vectashield. This was confusing since the same samples showed positive immunofluorescence labelling in PBS, suggesting a quenching of AF647 ON state in Vectashield. To understand this, we performed a quantitative analysis of the AF647 fluorescence intensity in Vectashield. Finally, based on our quantitative analysis, we could optimize imaging conditions in Vectashield to obtain 3D dSTORM images of targets, such as neurofilaments in neuronal cell line and axonal initial segments in primary mouse neurons.

## 2. Results

The aim of our study was to image neuronal proteins in Vectashield with dSTORM. To achieve this, we first established dSTORM imaging of microtubules in Vectashield (**Supplementary Fig. 1**), as described in the literature. Surprisingly, however, dSTORM imaging of neurofilament light chain (NfL) in neuronal ND7/23 cell line and axonal initial segments in primary mouse cortical neurons at first seemed not to be possible. The reason for this was a substantial loss of AF647 fluorescence in Vectashield compared to phosphate-buffered saline (PBS) (**Fig. 1a, c, d**). Since we used a secondary antibody conjugated to a new variant of AF647, AF Plus 647 (AF647+), we did a control experiment in which we compared conventional AF647 and AF647+. This experiment showed that fluorescence intensity of both AF647 and AF647+ decreases after addition of Vectashield (**Supplementary Fig. 2**). Consequently, the cells labelled with AF647 or AF647+ look completely dark in Vectashield. Since dyes are quenched in the fluorescent ON state, it is very difficult to distinguish labelled cells from the background. To exclude a possibility of antibody washout associated with imaging media change, we performed a control experiment with PBS. This experiment showed no decrease of fluorescence intensity (**Supplementary Fig. 3**). As an additional control, we show that Vectashield does not quench Alexa Fluor 488 Plus (AF488+; **Fig. 1b**) or other orange and red dyes, such as AF555 and AF633 (**Supplementary Fig. 4**).

**Figure 1.**
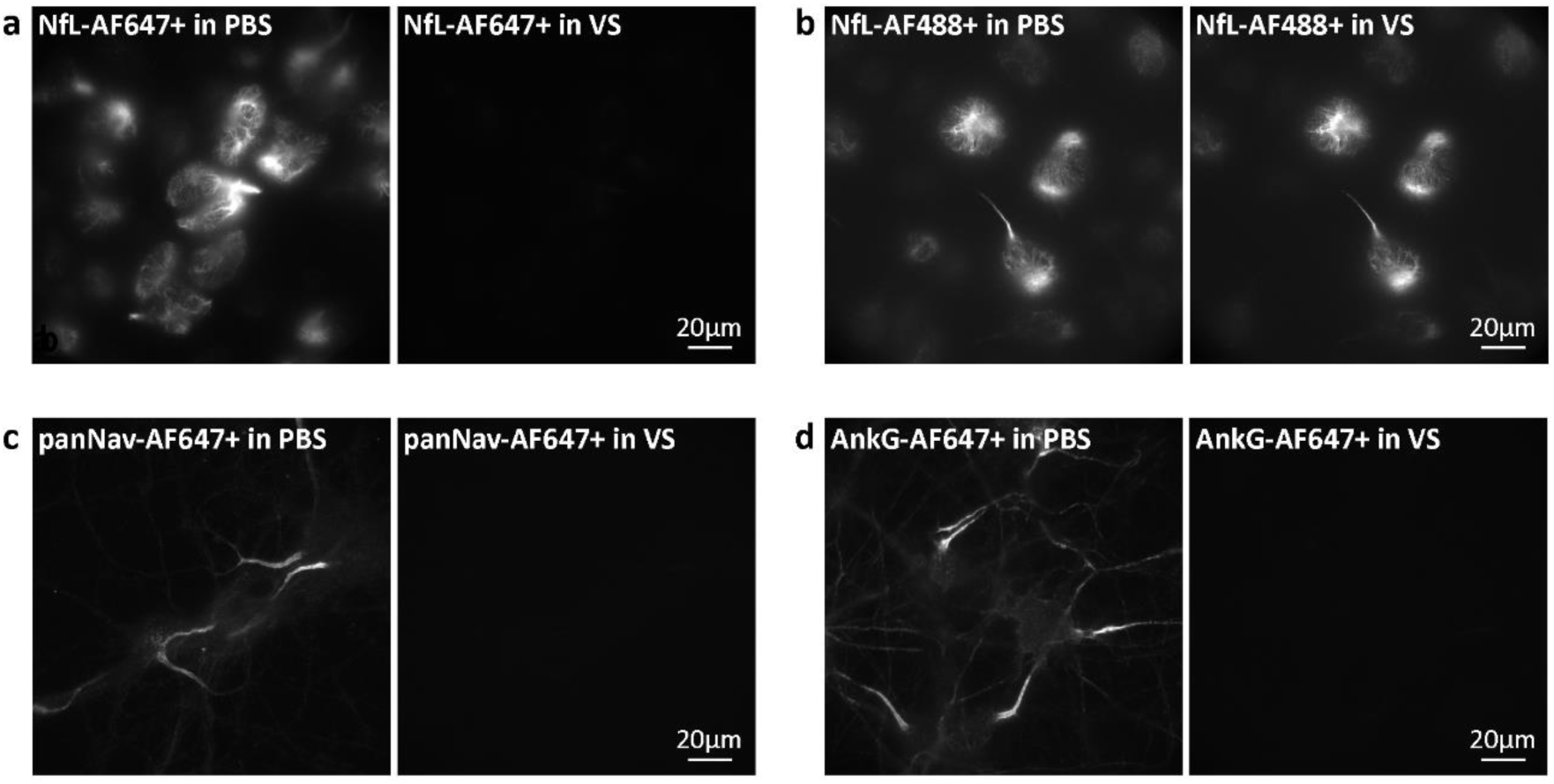
Effect of Vectashield (VS) on AF647+ and AF488+ fluorescence intensity. Widefield images of (a) AF647+ labelled neurofilament light chain (NfL) in ND7/23 cells, (b) AF488+ labelled NfL in ND7/23 cells, (c) AF647+ labelled sodium channels (panNa_v_) in mouse cortical neurons (MCN), (d) AF647+ labelled ankG in MCN. In (a-d), the same field of view is shown in PBS and in VS. Brightness and contrast are linearly adjusted to show the same display range, so that the effects of two imaging media can be compared.

To quantify Vectashield-induced fluorescence intensity changes, ND7/23 cells were transfected with a plasmid containing nuclear localization sequence (NLS)-mCherry fusion protein (**Fig. 2a**). We chose to use NLS-mCherry for intensity measurements for multiple reasons. First, it is uniformly distributed within the nucleus, and can be easily labelled with the same primary antibody, followed by either AF647 or AF488-conjugated secondary antibody. Additionally, mCherry gives a bright fluorescent signal in 561 (mCherry) channel that is not affected by any of the imaging media. This can be used for reliable identification of labelled regions of interest and intensity measurements of AF647 and AF488 before and after imaging media change (**Fig. 2b; Supplementary Fig. 5**). Quantitative analysis of fluorescence intensity changes shows that in Vectashield, there is an intensity drop to about 15 % of the initial value for AF647, and no loss of intensity for AF488 (**Fig. 2d**). We could also show that quenched AF647 signal can be recovered after a washing step **(Fig. 2c, d**).

**Figure 2.**
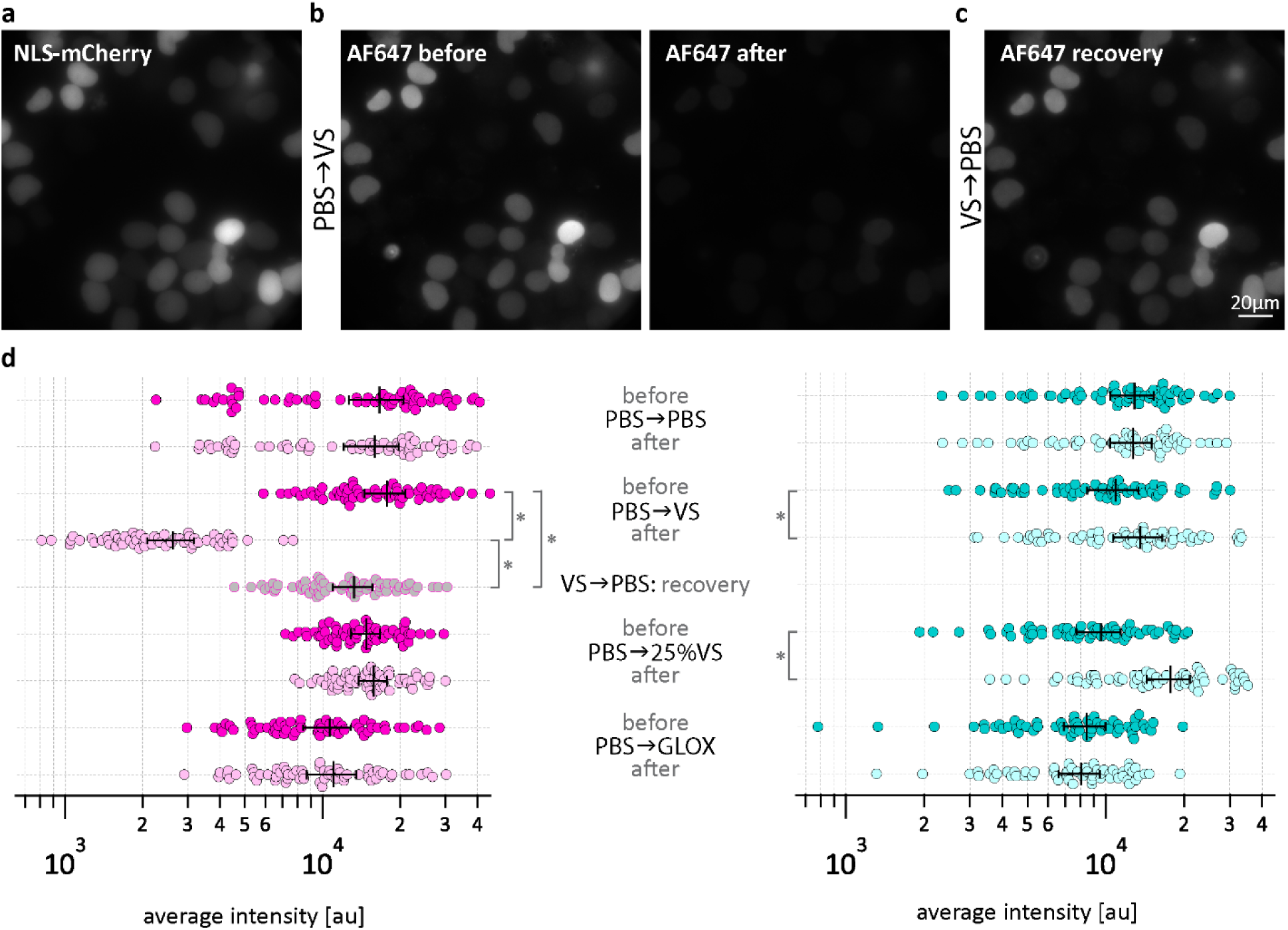
Quantification of fluorescence intensity changes of AF647 and AF488 in different imaging media. (a) Example widefield image of ND7/23 cells expressing NLS-mCherry. The cells are immunolabelled with anti-tRFP primary antibody, followed by AF647-conjugated secondary antibody. (b) Same field of view in 647 channel before and after PBS→Vectashield (VS) media change. (c) Same field of view in 647 channel in PBS 2.5h after removal of VS. (d) Dot plots show results of intensity change quantification in different imaging media for AF647 (magenta) and AF488 (cyan). In each case, cells were first imaged in PBS (before; darker shade on the graph), followed by medium change (after; lighter shade on the graph). Results of recovery experiment are shown in grey. Average fluorescence intensity values of all images from 3 experiments (20 images per medium were analysed in each experiment) are shown in the dot plot, including mean, and the standard error of the mean. Post-hoc Bonferroni comparisons (more information in Supplementary Details on Statistical Analysis) show that in all 3 experiments there is a significant decrease of AF647 intensity after changing the medium from PBS to Vectashield (PBS→VS; p<0.05), as well as significant increase after washing (VS→PBS: recovery; p<0.05). For AF488, there is a significant increase of intensity in VS and 25%VS (PBS→VS, p<0.05; PBS→25% VS, p<0.05). Average intensity (X-axis) is shown on a logarithmic scale.

To understand why the Vectashield-induced quenching was not reported earlier, we looked at the structures that were previously successfully imaged with dSTORM in Vectashield, such as tubulin and actin. Conventional widefield microscopy shows that AF647 is also quenched in the cells labelled with an AF647-conjugated tubulin antibody (**Fig. 3**) or AF647-phalloidin (**Supplementary Fig. 6**). However, AF647 quenching is less noticeable when looking at these structures compared to NfL or axonal initial segments. Despite the quenching, positively labelled microtubules or actin can easily be identified with widefield microscopy. On the other hand, this is not the case with other proteins, such as NfL in neuronal cell lines or components of axonal initial segments, such as voltage-gated sodium channels (Na_v_) or ankyrin G, in primary neurons. To identify positively labelled NfL or axonal initial segments, laser illumination had to be used. Even with laser and after we adjusted the image display (brightness/contrast) to show very dim pixel values (auto scale look-up table), NfL (**Fig. 4**), Na_v_ (**Fig. 5a**) and ankyrin G (**Fig. 5c**) labelling quality seems to be very poor, especially when compared to the images in GLOX buffer **(Fig. 5 b, d**). In addition, very soon after switching to dSTORM mode, and even under very low laser illumination, we noticed that NfL, Na_v_ and ankyrin G samples started blinking. This makes it challenging to search for the positive cells and regions of interest and to set the correct focus for 3D dSTORM. However, since blinking is a prerequisite for dSTORM, this prompted us to try to perform dSTORM.

**Figure 3.**
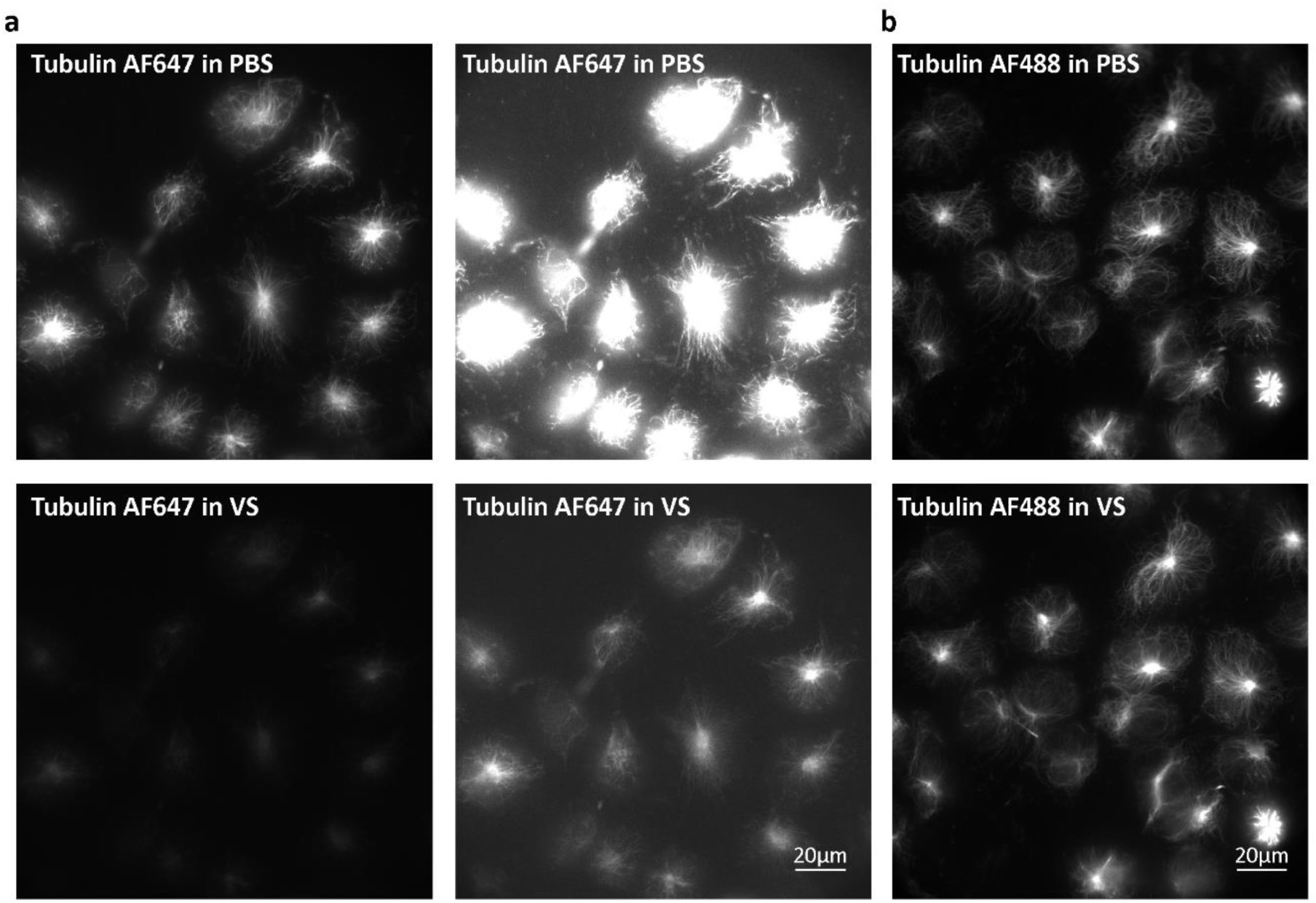
Vectashield (VS) effect on AF647 and AF488 labelled microtubules. (a) Widefield images of ND7/23 cells labelled with AF647-conjugated anti-Tubulin β3 (TUBB3) antibody in PBS (upper panels) and in VS (lower panels). To make the quenched AF647 signal in VS visible, same images with linearly adjusted brightness and contrast levels are shown on the right. (b) Widefield images of ND7/23 cells labelled with AF488-conjugated anti-Tubulin β3 (TUBB3) antibody in PBS (upper panel) and in VS (lower panel).

**Figure 4.**
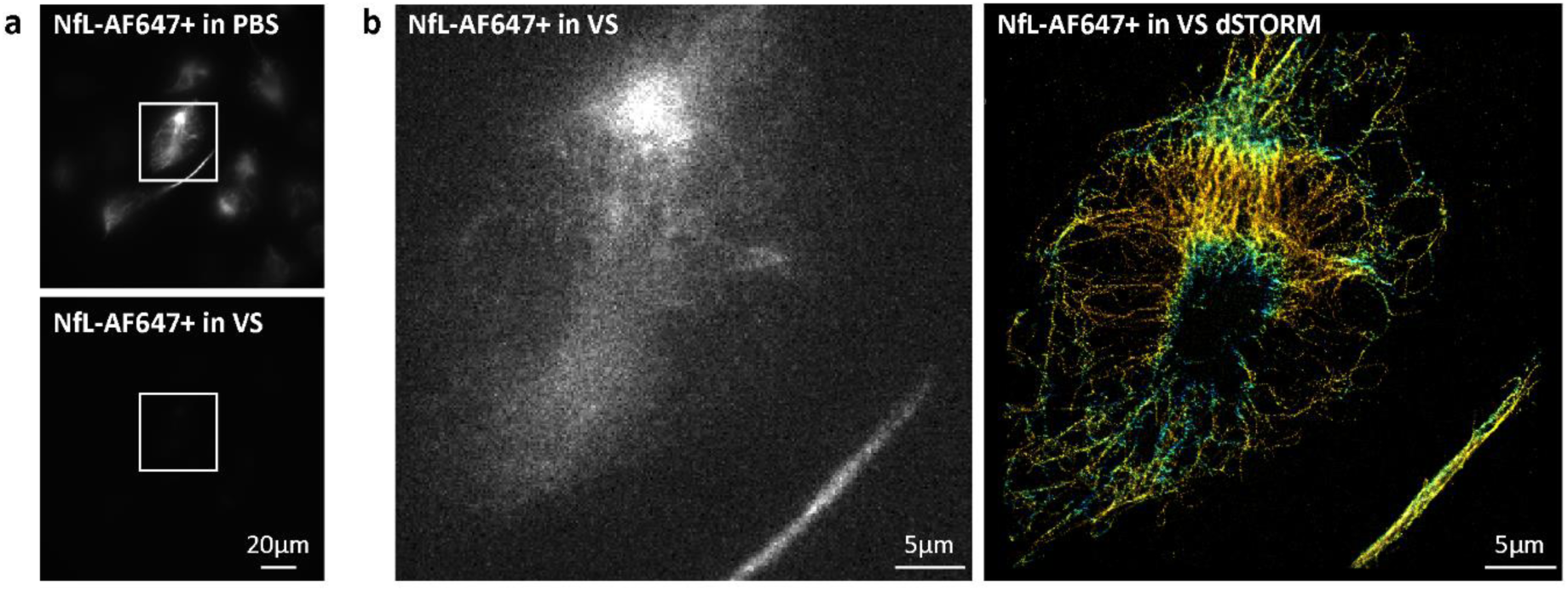
3D dSTORM super-resolution imaging of neurofilament light chain (NfL) in Vectashield (VS). (a) Widefield images of ND7/23 cells immunolabelled with anti-NfL primary antibody, followed by AF647+ conjugated secondary antibody. The same field of view is shown in PBS and in VS. Brightness and contrast are linearly adjusted to show the same display range, so that the effects of two imaging media can be compared. (b) TIRF image of the boxed region from panels in a. Image was acquired in VS, with 647 laser illumination prior to dSTORM imaging. To make the quenched AF647+ labelled cell visible, auto scale look-up table had to be used. Corresponding 3D dSTORM image is shown on the right panel.

**Figure 5.**
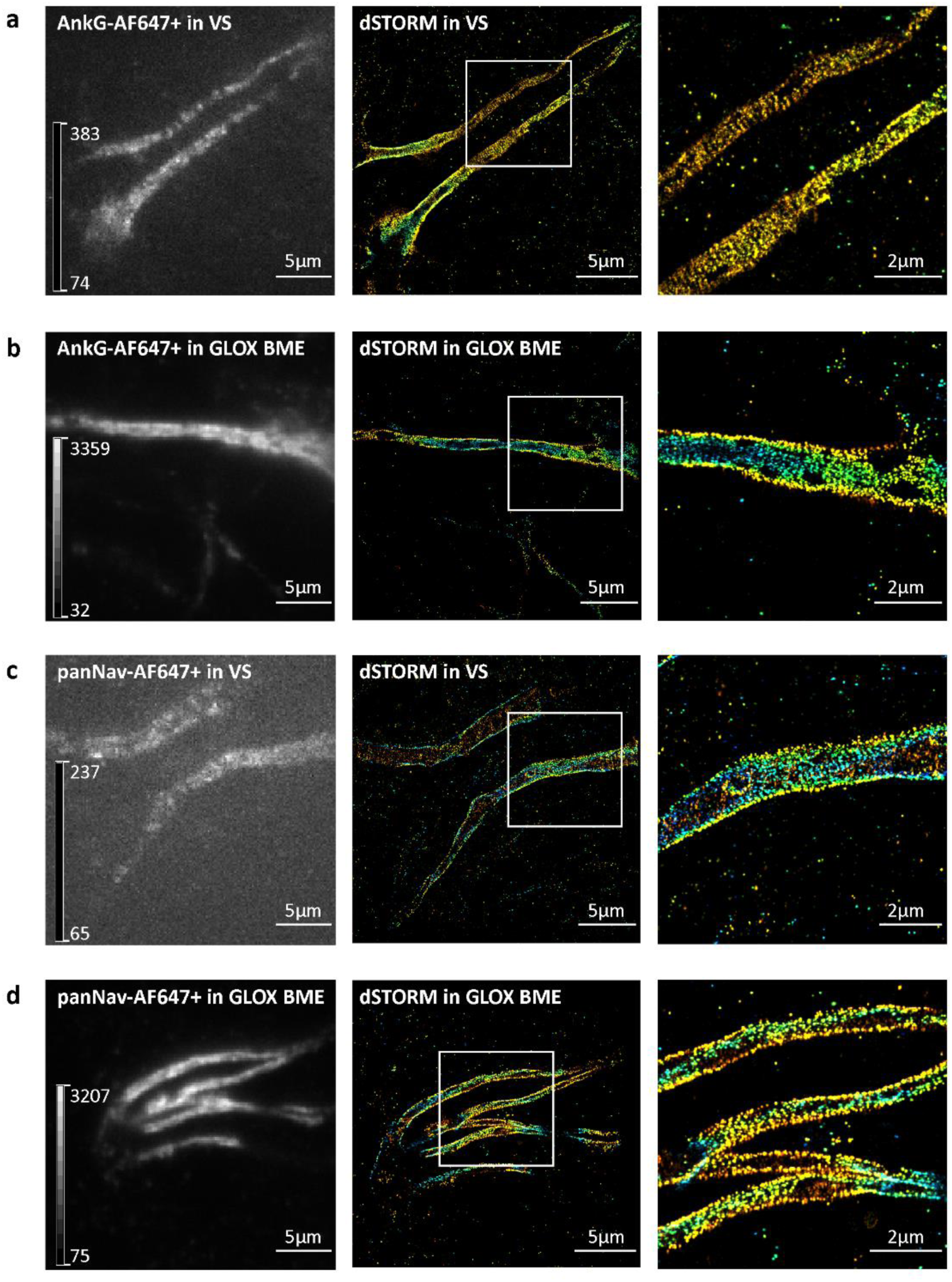
3D dSTORM super-resolution imaging of axonal initial segments in Vectashield (VS) and GLOX BME. (a,b) Mouse cortical neurons (MCN) labelled with anti-ankyrin G (ankG) antibody. (c,d) MCN labelled with anti-sodium channel (panNa_v_) antibody. Left panels show TIRF images, acquired with 647 laser illumination prior to 3D dSTORM imaging. To make the quenched AF647+ in axonal initial segments in VS visible, autoscale look-up table (LUT) was used. LUT intensity bars show minimum and maximum grey values. Middle panels show corresponding 3D dSTORM images of ankG (a,b) and panNa_v_ (c,d). Right panels show magnification of the boxed regions in middle panels. Despite the quenching effect of VS, periodic pattern of ankG and sodium channels can still be resolved.

For dSTORM imaging, we first identified AF647+ labelled NfL in PBS with conventional fluorescence microscopy. In the next step, we changed to Vectashield and laser illumination. In Vectashield, 3D dSTORM was performed and we obtained SRM images with a resolution of less than 40 nm (**Fig. 4**), as estimated by Fourier ring correlation (FRC)^19^. However, this was only possible by identifying the labelled cell in PBS before adding Vectashield, i.e. before the image turned dark. The procedure of identifying the cells in PBS first and then switching to Vectashield for 3D dSTORM imaging also worked for the components of axonal initial segment labelled with AF647+, such as ankyrin G and Na_v_ (**Fig. 5**). However, the procedure of identifying the cells in PBS, and then changing from PBS to Vectashield, makes dSTORM in Vectashield less optimal than dSTORM in GLOX buffer.

In addition to identifying the labelled cells in PBS first, and then switching to Vectashield for dSTORM, imaging could be done in 25% Vectashield. Our quantitative analysis shows that 25% Vectashield does not induce quenching of AF647, similar to GLOX (**Fig. 2d**). This is important, since it allows us to identify labelled cells, evaluate the labelling quality and to pick brightly labelled cells, which is a prerequisite for successful dSTORM. However, this was only possible with AF647-labelled samples. As a comparison of AF647+ and AF647-labelled axonal initial segments (**Fig. 6)** and NfL (**Supplementary Fig. 7**) showed, AF647+ was quenched in 25% Vectashield, while AF647 was not. dSTORM imaging of AF647+ in 25% Vectashield was still possible, but the positively labelled cells had to be first identified in PBS (**Fig. 6**).

**Figure 6.**
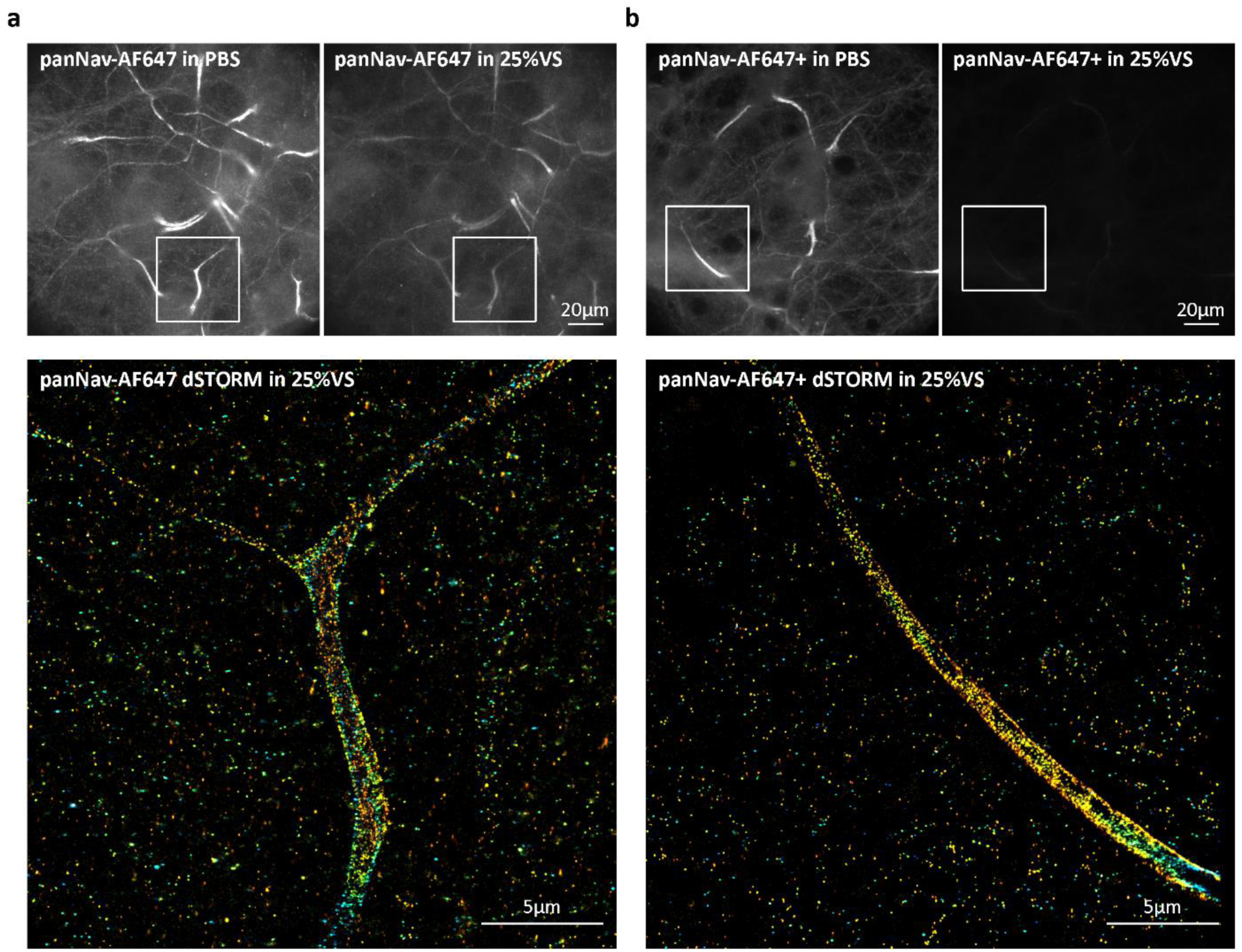
Comparison of 25% Vectashield (25%VS) effect on AF647 and AF647+ fluorescence. (a) Mouse cortical neurons (MCN) immunolabelled with anti-sodium channels (panNa_v_) antibody, followed by AF647-conjugated secondary antibody. (b) MCN immunolabelled with panNa_v_, followed by AF647+ conjugated secondary antibody. Upper panels show widefield images of the same field of view acquired in PBS and in 25%VS. Lower panels show corresponding 3D dSTORM images of the boxed regions from panels a and b.

## 3. Discussion

We show that one of the most common mounting media, Vectashield, induces quenching of AF647 and its improved variant, AF647+. As a consequence, the fluorescence intensity of both AF647 and AF647+ is lower in Vectashield. On the contrary, AF488 fluorescence is increased in both pure and 25% Vectashield (respectively to 125 % and 185 % of the initial PBS values). As our results show, diluting Vectashield in Tris-glycerol (25% Vectashield) overcomes the quenching problem for AF647, but not for AF647+. Our findings have important consequences for any type of fluorescence microscopy. AF647 is one of the most commonly used red dyes in super-resolution microscopy. In combination with Vectashield, it has been used for both dSTORM and SIM^12-14,20^. However, our results suggest that AF647 and AF647+ in Vectashield are not the optimal combination for any type of fluorescence microscopy, because the dyes get quenched. This is especially relevant for immunohistochemical staining because Vectashield is frequently used to mount tissue sections for confocal microscopy. If it is necessary to use AF647, 25% Vectashield is a good alternative. For AF647+, a different mounting medium needs to be used. Alternatively, by testing further dilutions of Vectashield, it might be possible to find an optimal concentration which would not induce AF647+ quenching.

Even though we only quantified effects of Vectashield and its dilutions on AF647 and AF488 fluorescence, similar changes might be happening with other fluorophores, other mounting media or other dSTORM imaging buffers. This is the reason why for any type of labelling, we would recommend to always first check the cells in buffers such as PBS. This is the only way to distinguish between failed immunocytochemistry/immunohistochemistry staining and fluorescence quenching phenomena. While Oliver and colleagues mention that some dyes are not stable in Vectashield (e.g. Cy2), they do not discuss any potential problems with stability of AF647 in Vectashield^13^. It was also suggested that Vectashield can induce cleavage of cyanine dyes and their derivatives. To test the hypothesis of Vectashield inducing cleavage of AF647, we performed a recovery experiment where we washed away Vectashield and after some time imaged the cells again in PBS. A quantitative analysis shows that the loss of AF647 fluorescence is reversible since it can be partially recovered. The recovery would not have been possible if the dye was cleaved. Since the dyes are affected in the ON state, Vectashield might be causing changes in the quantum yield, which on the other hand affects brightness of the dye.

Despite the quenching of fluorescence intensity in the ON state, AF647 dye molecules alternate between the ON and OFF states in Vectashield. Because of this, it is possible to perform dSTORM imaging in Vectashield. However, quenching of the fluorophores in the ON state hinders dSTORM imaging. One advantage of dSTORM over some of the other single molecule localizing techniques (e.g. PALM with photoactivatable proteins) is that it uses conventional labelling and starts with the fluorophores in the ON state^21^. This enables identification of brightly labelled cells and structures of interest. Because of quenching, this is not possible with Vectashield. In Vectashield, non-labelled (or poorly labelled cells) and quenched cells cannot be distinguished. Not being able to identify positively labelled cells is less critical for immunostainings of microtubules or actin filaments, which are abundant in the cells. Such immunostainings usually give a homogenous signal and almost any cell can be imaged with dSTORM. The cells can even be identified with transmitted light without the need for any fluorescent signal. However, this is not possible with most of the other targets and with highly polarized cells such as neurons. One example for this is axonal initial segment that we imaged in our manuscript. This complex structure has very unique molecular organisation^22,23^ and it cannot be identified based on neuronal morphology and transmitted light microscopy. Under a light microscope, axonal initial segment can only be identified by labelling its specific components and identifying them with fluorescence microscopy. To image such structures with dSTORM, it is necessary to identify the positively labelled cells with conventional fluorescence prior to dSTORM imaging. For this, the dye molecules need to be in the ON state. As this is not possible in Vectashield, we identified the AF647-labelled targets (NfL or axonal initial segments) in PBS and switched to Vectashield for dSTORM imaging. As an alternative, 25% Vectashield can be used for AF647. According to Olivier and colleagues, diluting Vectashield in Tris-glycerol (25% Vectashield) helps with reducing Vectashield’s autofluorescent background^13^. Furthermore, our analysis shows that 25% Vectashield does not induce quenching, which on its own makes it more suitable for imaging of AF647. However, according to our results, 25% Vectashield still quenched AF647+. Most of the studies until now used 25% Vectashield (or other dilutions) and AF647. This might also explain why Vectashield-induced quenching was not reported before. Our results show that even though both AF647 and AF647+ get quenched in Vectashield, they do not behave the same in 25% Vectashield. Because of this, additional care is necessary when choosing AF647-conjugated antibodies. In our experiments and as also reported by the manufacturer, AF647+ gives higher signal-to-noise and we would recommend it for dSTORM imaging.

Based on our imaging protocol, both AF647 and AF647+ can be used for dSTORM in Vectashield. However, depending on the target, imaging in Vectashield is more or less optimal, especially when compared to GLOX. It is important to emphasize that imaging in Vectashield was not a problem for structures such as microtubules, but became obvious when imaging other proteins, such as NfL in neuronal cell lines or voltage gated sodium channels and ankyrin G in axonal initial segments in primary neurons. This is probably a consequence of a relatively higher starting labelling intensity of microtubules. For example, anti-tubulin immunocytochemistry still shows relatively nicely labelled cells in Vectashield. Without seeing the image in PBS (before the addition of Vectashield), one might not even realize that there was quenching. Thus, depending on the target itself, the quality of the antibody and the labelling protocols, the quenching effect of Vectashield might be more or less noticeable. Until now, mainly cytoskeletal elements were imaged in Vectashield^14,16-18^. Since most of the biological structures are not as abundant as cytoskeletal elements, we predict that other targets might be affected similarly to the ones that we imaged in this manuscript. With our imaging protocol and quantitative analysis, it should be possible to optimize the imaging conditions in Vectashield. However, for such targets, it might be necessary to try additional dSTORM buffers, because Vectashield might not always be the most optimal choice.

## Materials and Methods

### Cell culture

Mouse neuroblastoma x rat neuron hybrid ND7/23 cells (ECACC 92090903, Sigma Aldrich) were grown in high glucose Dulbecco’s Modified Eagle Medium (DMEM; Thermo Fisher Scientific, cat. no. 41965062) supplemented with 10 % heat-inactivated fetal bovine serum (FBS; Thermo Fisher Scientific, cat. no. 10270106), 1 % penicillin-streptomycin (Sigma Aldrich, cat. no. P0781), 1 % sodium pyruvate (Thermo Fisher Scientific, cat. no. 11360039) and 1 % L-glutamine (Thermo Fisher Scientific, cat. no. 25030024) at 37 °C, 5 % CO2. FBS was heat-inactivated by incubation at 56 °C for 30 minutes (min). For more details, see Supplementary Information.

Primary mouse cortical neurons (MCN) from C57BL/6 embryonic day-17 mice were obtained from Thermo Fisher Scientific (cat. no. A15586). Neurons were thawed following the manufacturer’s recommendations and seeded in eight-well Lab-Tek II chambered #1.5 German coverglass (Thermo Fisher Scientific, cat. no. 155409). Neurons were maintained in B-27™ Plus Neuronal Culture System (Thermo Fisher Scientific, cat. no. A3653401), supplemented with 1 % penicillin-streptomycin (Sigma Aldrich, cat. no. P0781). Half of the culturing medium was changed twice per week.

### Constructs and cloning

NLS (nuclear localization sequence)-mCherry construct was made by inserting NLS-mCherry sequence into a commercially available pcDNA™3.1/Zeo(+) mammalian expression vector (Invitrogen). NLS sequence was added upstream of the mCherry gene (gift from Edward Lemke’s laboratory, EMBL, Heidelberg) by polymerase chain reaction (PCR). The following primers were used for cloning: 5’-GCTGGCGCTAGCACCATG**CCGCCGAAAAAAAAACGCAAAGTGGAAGATAGC**GTGAGCAAGGGCGAGGAG G-3’ (forward primer with NheI restriction site; NLS sequence is shown in bold) and 5’-CGCGCAGCGGCCGCTCACTTGTACAGCTCGTCCATGCCG-3’ (reverse primer with NotI restriction site). Generated plasmid sequence was confirmed by sequencing.

### Transfections (for AF647 and AF488 intensity measurements)

For Alexa Fluor 647 (AF647) and Alexa Fluor 488 (AF488) intensity measurement experiments, ND7/23 cells were transfected with NLS-mCherry plasmid. Cells were transfected using JetPrime (Polyplus-transfection, cat. no. 114-15) one day after seeding, at 80-85 % confluence, according to the manufacturer’s instructions. On the following day, NLS-mCherry expression was confirmed on an epifluorescent inverted microscope (Olympus CKX41) and immunostaining was performed.

### Immunocytochemistry stainings

For immunocytochemistry (ICC) stainings, ND7/23 cells were washed briefly with 0.01 M phosphate-buffered saline (PBS; 137 mM NaCl, 10 mM Na2HPO4, 1.8 mM KH2PO4, 2.7 mM KCl, pH 7.4). Tris-buffered saline (TBS; 20 mM Tris, 150 mM NaCl, pH 7.6) was used instead of PBS for immunostaining of neurofilament light chain (NfL) because of potential influence of PBS on phosphorylated neurofilaments. Afterwards, cells were fixed with 2 % paraformaldehyde (PFA; Sigma Aldrich, cat. no. 158127) at RT. For labelling of tubulin, cells were fixed as described previously^13^.

After fixation, cells were briefly washed again, blocked and permeabilized. Following primary antibodies were used: anti-tRFP (turbo red fluorescent protein) antibody (Evrogene, cat. no. AB233), mouse anti-neurofilament 70 kDa antibody, clone DA2 (Merck Millipore, cat. no. MAB1615), AF647-conjugated anti-Tubulin β3 (TUBB3) antibody (BioLegend, cat. no. 801210), AF488-conjugated anti-TUBB3 antibody (BioLegend, cat. no. 801203), anti-TUBB3 antibody (BioLegend, cat. no. 801202). For actin labelling, cells were incubated with phalloidin AF647 (Thermo Fisher Scientific, cat. no. A22287).

Following secondary antibodies were used: goat-anti rabbit AF647 (Thermo Fisher Scientific, cat. no. A-21245), goat anti-rabbit AF488 (Thermo Fisher Scientific, cat. no. A-11034), goat anti-mouse AF647 Plus (Thermo Fisher Scientific, cat. no. A32728), goat anti-mouse AF488 Plus (Thermo Fisher Scientific, cat. no. A32723), goat anti-mouse AF647 (Thermo Fisher Scientific, cat. no. A21236), goat anti-mouse AF555 (Thermo Fisher Scientific, cat. no. A21424), goat anti-mouse AF633 (Thermo Fisher Scientific, cat. no. A21052).

MCN were fixed from 15th to 19th day *in vitro* (DIV) with 4 % EM grade PFA (Electron Microscopy Sciences, cat. no. 15710) diluted in PEM buffer (80 mM PIPES, 2 mM MgCl_2_, 5 mM EGTA, pH 6.8) for 15 minutes at RT. After fixation, background fluorescence was quenched with sodium borohydride (Sigma Aldrich, cat. no. 71320), cells were washed 3 times (10 minutes each wash) with PBS, blocked and permeabilized. Following primary antibodies were used: mouse monoclonal anti-pan sodium channel antibody (panNa_v_; Sigma Aldrich, cat. no. S8809) and mouse monoclonal anti-ankyrin G antibody (Santa Cruz, cat. no. sc-12719). Goat anti-mouse secondary antibodies conjugated with AF647 Plus or AF647 were used.

Details on ICC staining steps and antibodies used in each figure are provided in Supplementary tables S1 and S2 and S3.

After labelling, ND7/23 cells and MCN were washed 3 times (5 minutes each wash) and imaged on the same day. Cells stained with anti-tRFP (NLS-mCherry transfected cells) were also imaged on the following day.

### Microscope configuration

Widefield epifluorescence and 3D dSTORM imaging were performed on an N-STORM 4.0 microscope from Nikon Instruments. More specifically, this is an inverted Nikon Eclipse Ti2-E microscope (Nikon Instruments), equipped with XY-motorized stage, Perfect Focus System, an oil-immersion objective (HP Apo TIRF 100×H, NA 1.49, Oil) and N-STORM module. Setup was controlled by NIS-Elements AR software (Nikon Instruments). Fluorescent light was filtered through following filter cubes: 488 (AHF; EX 482/18; DM R488; BA 525/45), 561 (AHF; EX 561/14; DM R561; BA 609/54), Cy5 (AHF; EX 628/40; DM660; BA 692/40) and Nikon Normal STORM cube (T660lpxr, ET705/72m). Filtered emitted light was imaged with ORCA-Flash 4.0 sCMOS camera (Hamamatsu Photonics). For epifluorescent widefield imaging, fluorescent lamp (Lumencor Sola SE II) was used as a light source. For 3D dSTORM imaging of neurofilaments, 647 nm laser (LU-NV Series Laser Unit) was used and cylindrical lens was introduced in the light path^11^.

### Imaging of tRFP labelled cells (for AF647 and AF488 intensity measurements)

tRFP labelled cells were first briefly checked in PBS using brightfield illumination. For each well in the Lab-Tek we did following: picked randomly 30 fields of view (stage positions) using brightfield illumination, saved xyz coordinates of each field of view in NIS-Elements AR software and acquired images automatically by using NIS-Elements ND multipoint acquisition module, which allowed us to go to the same position each time. The images were acquired in widefield mode, with 10 ms exposure time, 1024×1024 pixels frame size and 16-bit image depth. To provide that the cells were always properly focused, we used an autofocusing function of ND multipoint acquisition module.

Imaging was done first in PBS, using 561 (mCherry channel) and either 488 (AF488 channel) or Cy5 (AF647 channel) filter cube, depending on the labelling condition. Excitation light intensity for mCherry and AF647 channels was 10 % and for AF488 channel 5 %. Afterwards, PBS was replaced with one of the following imaging media: PBS, 100% Vectashield (Biozol, cat. no. VEC-H-1000), 25% Vectashield or GLOX. More details on imaging media composition can be found in Supplementary Information. For AF647 recovery experiments, 100% Vectashield was removed and cells were washed twice with PBS. After 2.5 h, cells were washed one more time with PBS and imaging was repeated. To provide enough data for analysis, each experiment was repeated at least three times for AF488 labelled cells and AF647 labelled cells.

### Image analysis and intensity measurements

Intensity measurements for quantitative analysis of AF647 and AF488 were done in Fiji/ImageJ^24^. Images in ND2 format were opened using Bio-Formats plugin^25^ and converted to tiff before the analysis. Original bit depth (16-bit) was used for analysis. For presentation purposes, brightness and contrast of 16-bit images were linearly adjusted in Fiji, images were converted to 8-bit and imported in Adobe Illustrator. For Figure 5, images were autoscaled and look-up table (LUT) intensity bars were added automatically, using NIS Elements software.

More details on image analysis and intensity measurements are provided in Supplementary Information.

### 3D dSTORM imaging

3D dSTORM (direct stochastic optical reconstruction microscopy) imaging was performed by using the N-STORM module of Nikon microscope described above. For imaging, oil-immersion objective (HP Apo TIRF 100×H, NA 1.49, Oil) and 647 nm laser (LU-NV Series Laser Unit) were used. Fluorescent light was filtered through a Nikon Normal STORM filter cube. Filtered emitted light was imaged with ORCA-Flash 4.0 sCMOS camera (Hamamatsu Photonics) with cylindrical lens introduced in the light path^11^.

Fluorescent and brightfield images of phalloidin AF647, NfL and tubulin labelled cells in PBS and Vectashield were acquired using fluorescent light source and Cy5 filter cube or 488 filter cube (more details are provided in Supplementary Information). After widefield acquisition, NfL and tubulin labelled cells were imaged with 3D dSTORM. Imaging was performed in total internal reflection fluorescence (TIRF) or highly inclined and laminated sheet microscopy (HiLo) mode with continuous 647 nm laser illumination (full power). Frame size was 256×256 pixels and image depth 16-bit. For each 3D dSTORM image, 30,000 frames were acquired at 33 Hz.

Imaging of axon initial segments was performed in the way described for NfL and tubulin, with some modifications. For imaging of sodium channels, 30,000 frames were acquired at 50 Hz or 33 Hz, and for imaging of ankyrin G, 20,000 or 30,000 frames were acquired at 25 Hz or 33 Hz. 3D STORM imaging was done in 100% Vectashield, 25% Vectashield or GLOX β-mercaptoethanol (GLOX BME).

Calibration of 3D dSTORM was done previously using TetraSpeck Microspheres (Thermo Fisher Scientific, cat. no. T7279) following NIS-Elements’ instructions. 3D dSTORM image processing was performed in NIS-Elements AR software. Molecule identification settings were set to defaults for 3D dSTORM analysis: minimum width 200 nm, maximum width 700 nm, initial fit width 300 nm, max axial ratio 2.5 and max displacement 1. Minimum height for peak detection was set to 200 (NfL), 750 (tubulin), 100 (Na_v_) and 500 (ankG), and localization analysis was performed with overlapping peaks algorithm. After reconstruction of NfL images, resulting molecule lists were exported as text files and analyzed by Fourier ring correlation (FRC)^19^ to determine the image resolution. Based on FRC resolution estimate (35-40 nm), we reconstructed 3D dSTORM images with Gaussian rendering size of 15 nm in NIS Elements AR. Brightness and contrast were adjusted with Gaussian rendering intensity and final 3D dSTORM image was exported as a tiff image. Z rejected molecules were excluded from resolution analysis and from final images.

### Statistical analysis

Statistical analyses (ANOVA and post-hoc analyses) were carried out with IBM SPSS Statistics Version 25, Armonk, New York, USA. The power analysis to determine the optimal sample size was carried out with the Statistical Tree Power Calculator, QFAB (Queensland Facility for Advanced Bioinformatics), Brisbane, Australia. Further details on statistical analysis are in Supplementary Information.

## Acknowledgements

This study is supported by the Emmy-Noether programme (project number 317530061 to INS) of the German Research Foundation and the Werner Reichardt Centre for Integrative Neuroscience (CIN) at the Eberhard Karls University of Tübingen. The CIN is an Excellence Cluster funded by the German Research Foundation within the framework of the Excellence Initiative (EXC 307).

## Author Contributions

AA and NS performed the experiments and image analysis, with the help of INS. RS performed the statistical analysis with the input from INS. INS designed the experiments with statistical advice from RS. INS wrote the manuscript with the input from AA, NS and RS. INS supervised the project.

## Data availability

Data available on request from the corresponding author.

## Competing Interests Statement

The authors declare no competing interests.

## Notes

#### Summary of Updates

dSTORM imaging data updated to include more targets, such as voltage-gated sodium channels and ankyrin G in primary mouse cortical neurons; comparison with GLOX imaging is also included

