## Supplementary Information for "Vectashield-induced fluorescence quenching hinders conventional and super resolution microscopy"

### Table of Content Supplementary Information


#### Supplementary Figures

**Supplementary Figure 1.** 3D dSTORM of AF647 and AF647+ labeled microtubules in Vectashield

**Supplementary Figure 2.** Comparison of Vectashiled (VS) effect on AF647+ and AF647 labeled ND7/23 cells and mouse cortical neurons (MCN)

**Supplementary Figure 3.** Control experiments with PBS to PBS media change show no effect on fluorescence intensity of AF647+, AF647 and AF488+ labeled neurofilaments

**Supplementary Figure 4.** Comparison of PBS vs. Vectashield effect on AF555 and AF633 labeled neurofilaments

**Supplementary Figure 5.** Additional examples of images used for quantification of intensity changes for AF647 and AF488

**Supplementary Figure 6.** Comparison of PBS vs. Vectashield effect on AF647-phalloidin

**Supplementary Figure 7**. Comparison of 25% Vectashield effect on AF647+ and AF647 labeled neurofilaments

#### Supplementary Tables

**Supplementary Table S1.** Detailed overview of imunocytochemistry staining steps in ND7/23 cell line

**Supplementary Table S2.** Detailed overview of imunocytochemistry staining steps in primary mouse cortical neurons

**Supplementary Table S3.** Summary table of antibodies used in the manuscript

#### Supplementary Material and Methods

Cell culture

Imaging of actin (phalloidin AF647), NfL and tubulin labelled cells

Imaging media for dSTORM

Image analysis and intensity measurements

#### Supplementary Details on Statistical Analysis

Preparation of data analysis based on pilot work

Power calculations, randomization and design

Deriving hypotheses based on prior observations

Data analyses

Separate ANOVA for AF647 in each of the 3 experiments

Analysis of AF647 recovery

Separate ANOVA for AF488 in each of the 3 experiments

#### Supplementary References

### Supplementary Figures

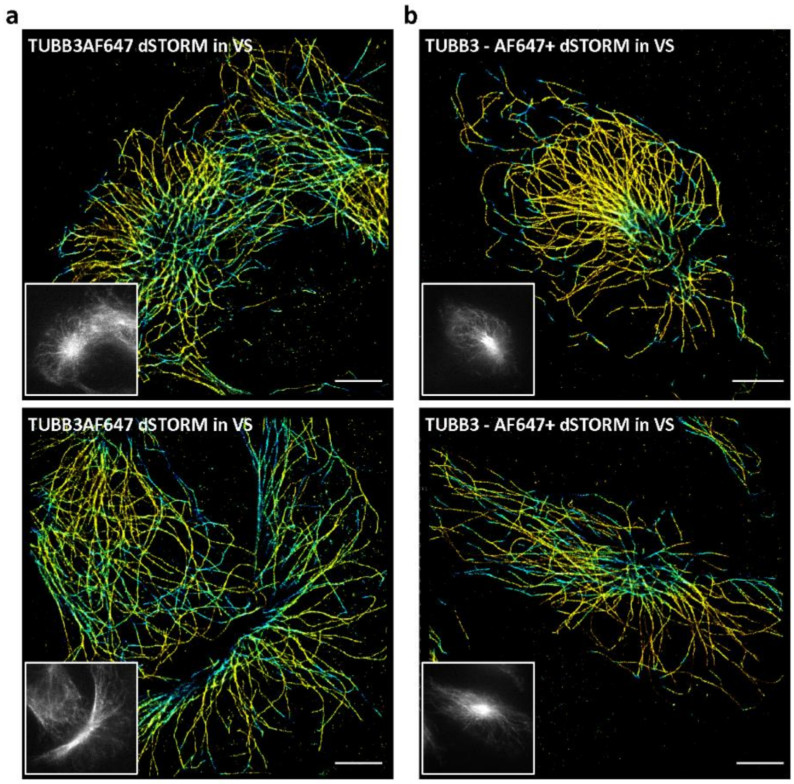


**Supplementary Figure 1.** 3D dSTORM of AF647 and AF647+ labelled microtubules in Vectashield (VS). 3D dSTORM images of ND7/23 cells immunolabelled with (a) AF647-conjugated anti-Tubulin β3 (TUBB3AF647) antibody or (b) anti-Tubulin β3 primary antibody, followed by AF647+ conjugated secondary antibody. Insets show TIRF images of the corresponding cells, acquired with 647 laser illumination prior to dSTORM imaging. For all the insets, auto scale look-up table is used. Scale bars: 5 µm.

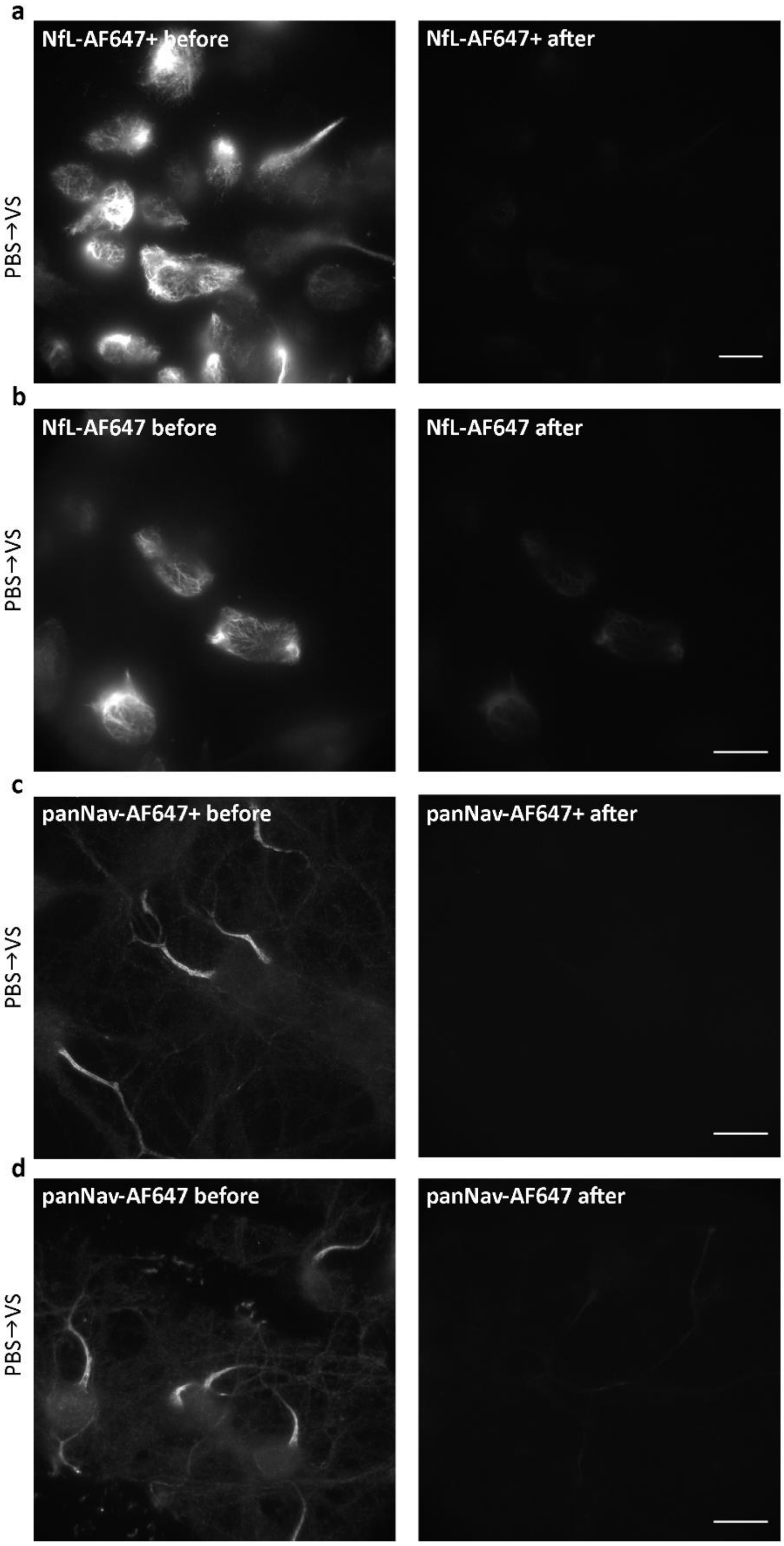


**Supplementary Figure 2.** Comparison of Vectashiled (VS) effect on AF647+ and AF647 labelled (a,b) ND7/23 cells and (b,d) mouse cortical neurons (MCN). ND7/23 cells labelled with anti-neurofilament light chain (NfL) primary antibody, followed by (a) AF647+ or (b) AF647 conjugated secondary antibody. MCN are labelled with anti-sodium channel (panNa$_V$) primary antibody, followed by AF647+ (c) or AF647 (d) conjugated secondary antibody. Left panels show widefield images of cells in PBS, before medium change and right panels show widefield images of cells in VS, after medium change. Brightness and contrast are linearly adjusted to show the same display range for images taken before and after media change. Scale bar: 20 µm.

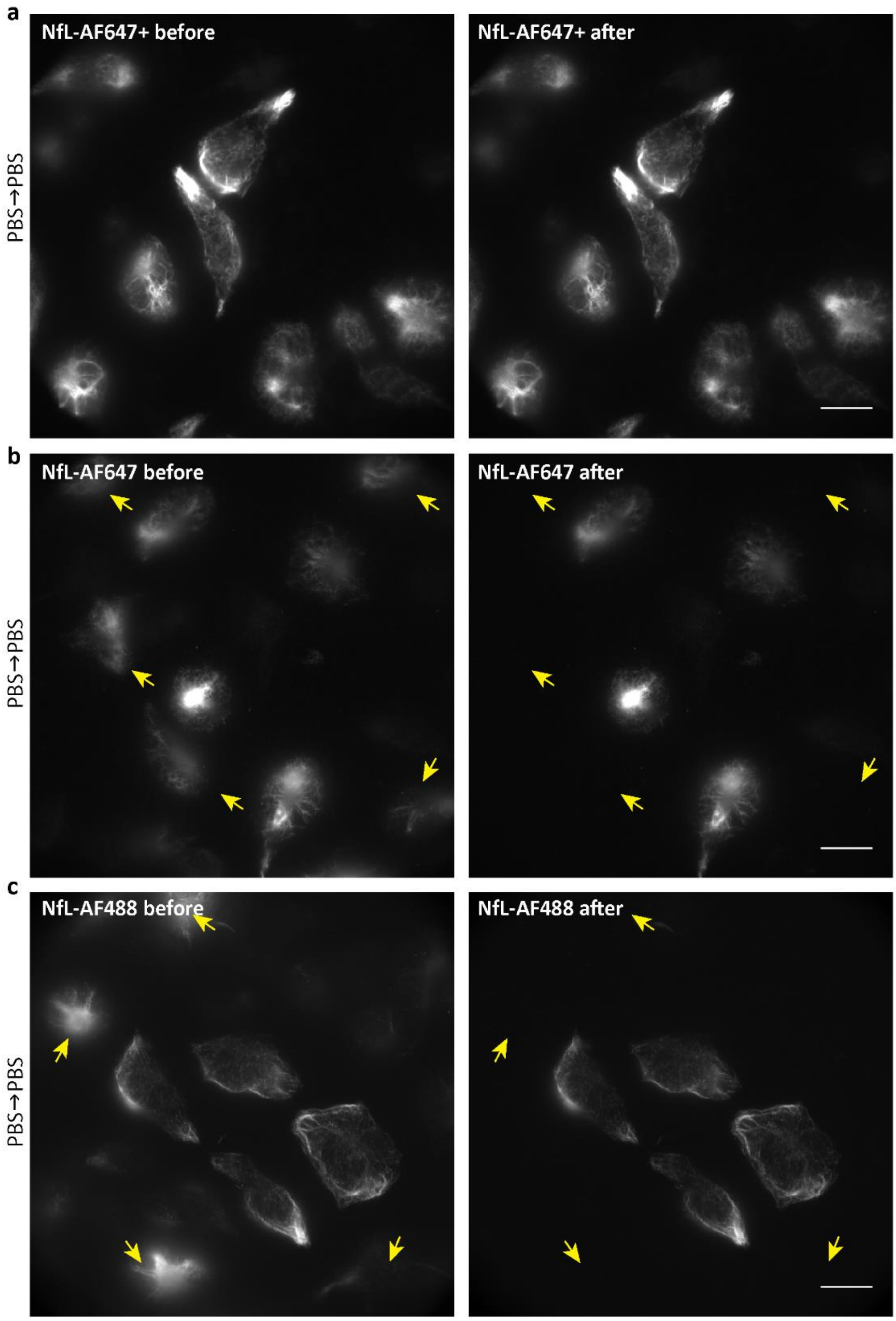


**Supplementary Figure 3.** Control experiments with PBS to PBS media change show no effect on fluorescence intensity of AF647+, AF647 and AF488+ labeled neurofilaments. ND7/23 cells labelled with anti-neurofilament light chain (NfL) primary antibody, followed by AF647+ (a), AF647 (b) or AF488+ (c) conjugated secondary antibody. Left panels show widefield images of cells in PBS before, and right panels show widefield images of cells in PBS after medium change. Please note that some cells got washed away during medium change – they cannot be identified in the image after medium change, and are marked with yellow arrows. For all the images brightness and contrast are linearly adjusted to show the same display range for images taken before and after media change. Scale bar: 20 µm (a,b,c).

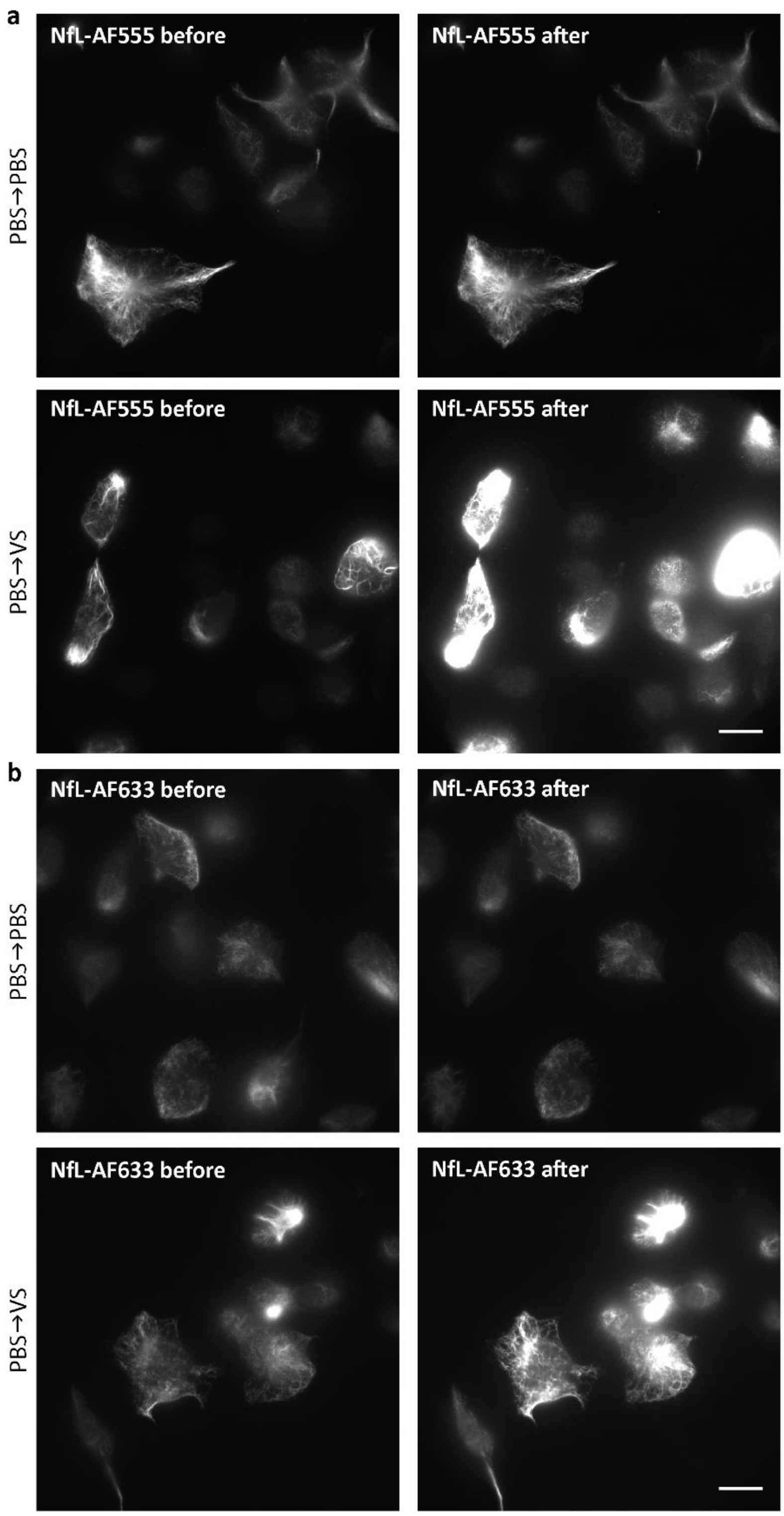


**Supplementary Figure 4.** Comparison of PBS vs. Vectashield effect on AF555 and AF633 labelled neurofilaments. ND7/23 cells labelled with anti-neurofilament light chain (NfL) primary antibody, followed by AF555- (a) or AF633-conjugated secondary antibody (b). Left panels show images before medium change in PBS and right panels show images after medium change in PBS (a and b, upper panels) or Vectashield (a,b, lower panels). For all the images brightness and contrast are linearly adjusted to show the same display range for images taken before and after media change. Scale bar: 20 µm (a,b).

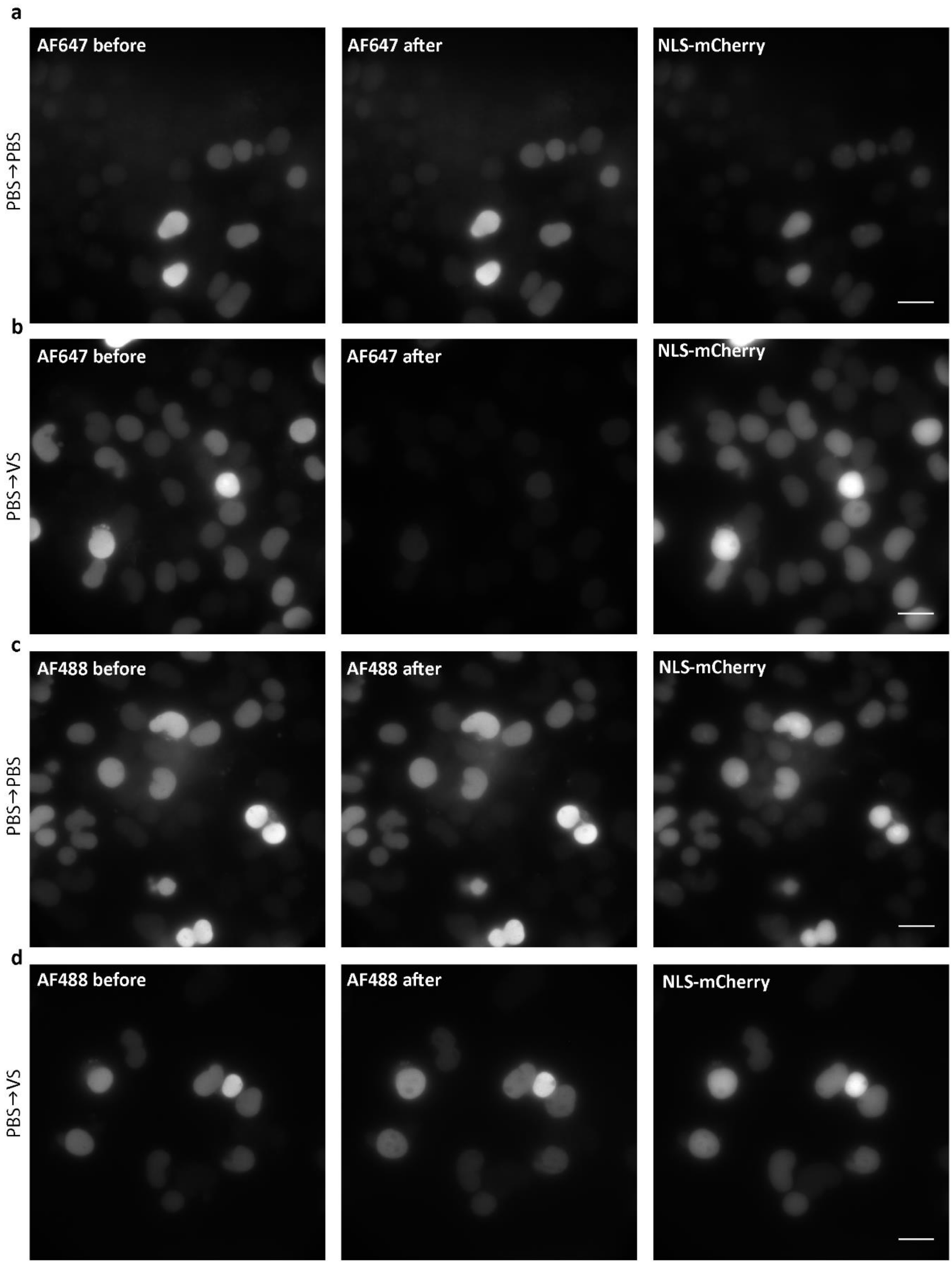


**Supplementary Figure 5.** Additional examples of images used for quantification of intensity changes for AF647 (a,b) and AF488 (c,d). Left panels show images before medium change (PBS) and middle panels show images after medium change (PBS or Vectashield). Right panels show corresponding mCherry channel images after medium change (PBS or Vectashield). In all of the panels, brightness and contrast are linearly adjusted to show the same display range for images taken before and after media change. Scale bar: 20 µm (a-d).

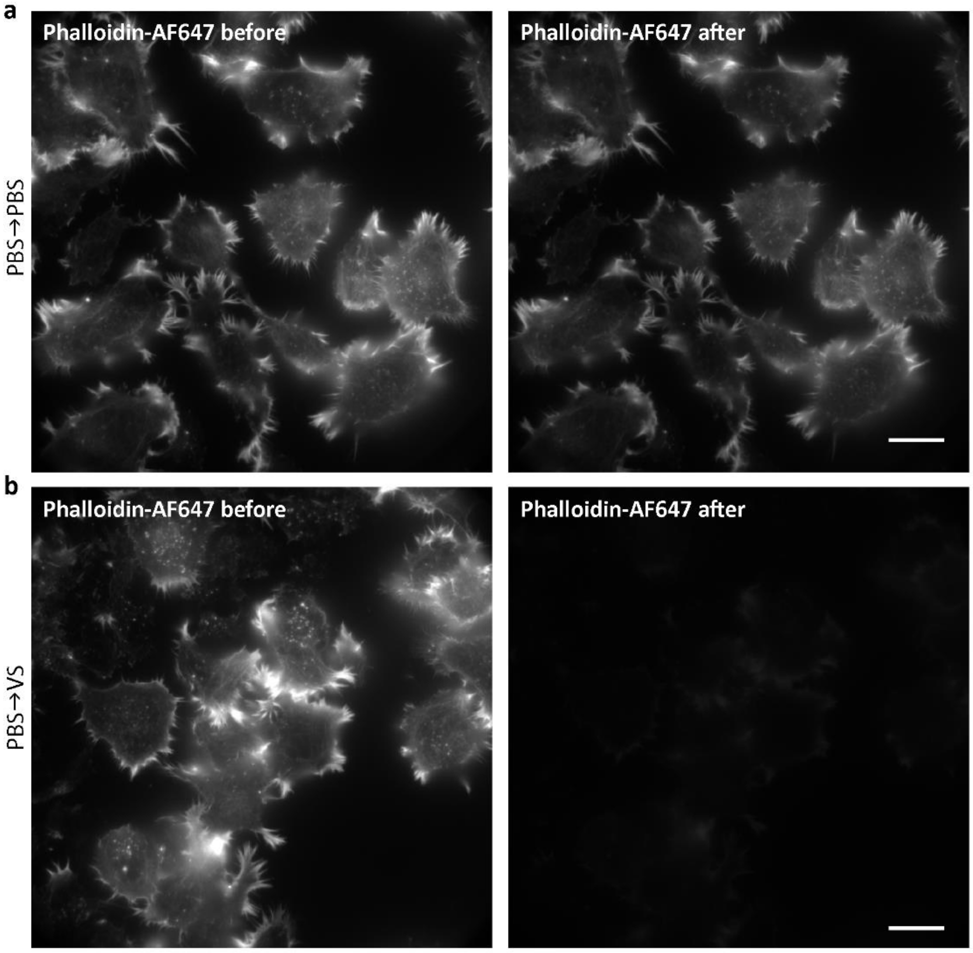


**Supplementary Figure 6.** Comparison of PBS vs. Vectashield effect on AF647-phalloidin. ND7/23 cells labelled with AF647 phalloidin. (a) Widefield images of cells in PBS before medium change (left panel) and after medium change (right panel). (b) Widefield images of cells in PBS (left panel) and in Vectashield (right panel). In a and b, brightness and contrast are linearly adjusted to show the same display range for images taken before and after media change. Scale bar: 20 µm (a,b).

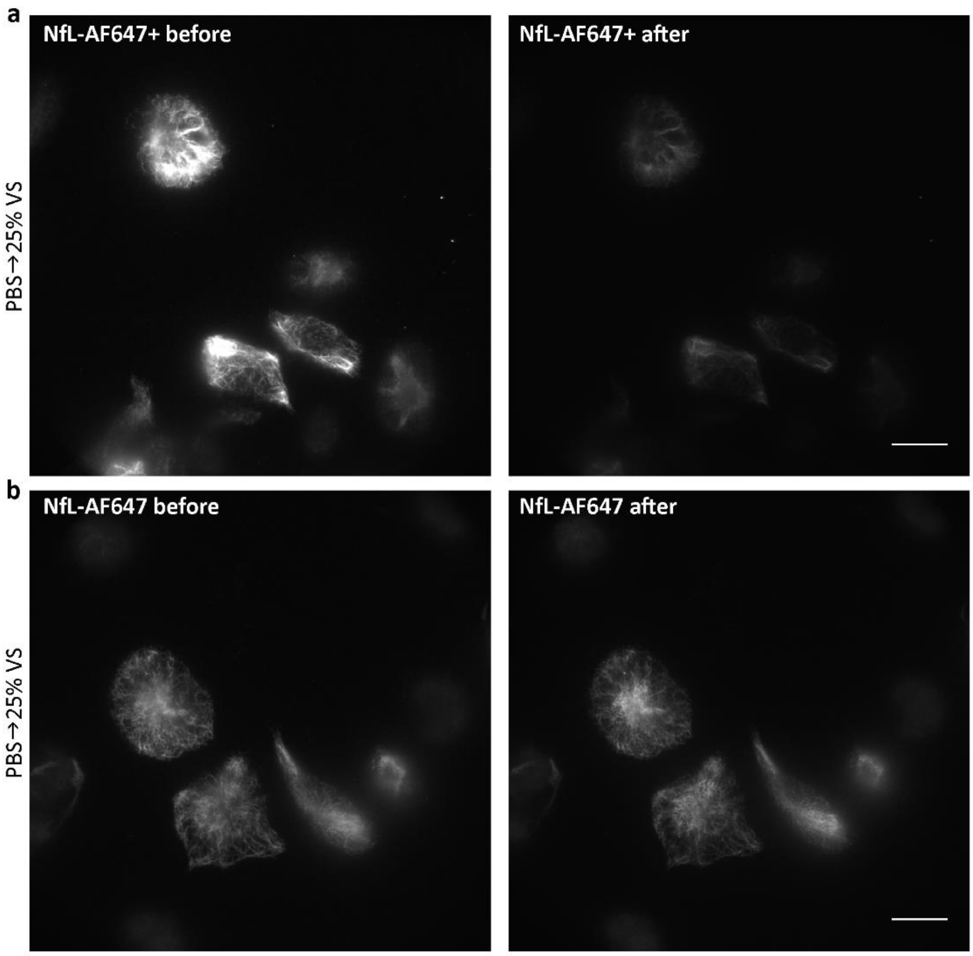


**Supplementary Figure 7**. Comparison of 25% Vectashield effect on AF647+ and AF647 labelled neurofilaments. ND7/23 cells labelled with anti-neurofilament light chain (NfL) primary antibody, followed by AF647+ (a) or AF647-conjugated secondary antibody (b). Widefield images of cells before medium change in PBS (left panels) and in after medium change in 25% Vectashield (right panels). In (a) and (b), brightness and contrast are linearly adjusted to show the same display range for images taken before and after media change. Scale bar: 20 µm (a,b).

### Supplementary Tables

**Supplementary Table S1. Detailed overview of imunocytochemistry staining steps in ND7/23 cell line**

| Target | NLS-mCherry | Neurofilament light chain (NfL) | Tubulin | Actin |
|---|---|---|---|---|
| **Fixation** | 2% PFA, 10min RT | 2% PFA, 15min RT | -20°C methanol for 10min | 2% PFA, 15min RT |
| **Blocking solution and duration** | 10% goat serum (GS)/0.5% Triton/PBS, 1h RT | 3% GS/0.3% Triton/TBS, 40min RT | 5% BSA/PBS, 1h RT | 5% BSA/0.3% Triton/PBS, 1.5h RT |
| **Primary antibody or dye** | Rabbit anti-tRFP (Evrogene, AB233) | Mouse anti-NfL 70 kDa, clone DA2 (Merck Millipore, MAB1615) | AF647 anti-TUBB3 (801210), AF488 anti-TUBB3 (801203) or mouse anti-TUBB3 (801202)** | AF647 phalloidin (A22287) |
| **Primary antibody dilution and incubation time** | 1:500 in 5% GS/0.1% Triton/PBS (at least 18h, 4°C) | 1:250 in 1% GS/0.3% Triton/TBS, 2.5h RT, followed by overnight incubation at 4°C | 1:100 in blocking serum, directly conjugated antibodies 3h RT, non-conjugated antibody ON at 4°C | 0.25µM/PBS, ON at 4°C, followed by 4h RT |
| **Washing steps** | All in 0.01M PBS | All in 0.01M TBS | All in 0.01M PBS | All in 0.01M PBS |
| **Secondary antibody*** | Goat anti-rabbit AF647 (A21245) or AF488 (A11034) | Goat anti-mouse AF647 Plus (A32728), AF488 Plus (A32723), AF647 (A21236), AF555 (A21424) or AF633 (A21052) | Goat anti-mouse AF647 Plus (A32728) | |
| **Secondary antibody dilution and incubation time** | 1:500 in 5% GS/0.1% Triton/PBS, 1h RT | 1:500 in 1% GS/0.3% Triton/TBS, 1h RT | 1:500 in blocking serum, 1h RT | |
| **Remarks** | | Used TBS instead of PBS | Extraction in 0.5% Triton/microtubule stabilization buffer, 10s, before fixation | |

*all secondary antibodies and AF647 phalloidin were purchased from Thermo Fisher. Product numbers are indicated in brackets.

**antibodies against tubulin were obtained from BioLegend. Product numbers are indicated in brackets.

**Supplementary Table S2. Detailed overview of imunocytochemistry staining steps in primary mouse cortical neurons**

| **Target** | **Sodium channels** | **Ankyrin G** |
|---|---|---|
| **Antibody** | Mouse anti-pan sodium channel (Sigma Aldrich, S8809) | Mouse anti-ankyrin G (Santa Cruz, 12719) |
| **Fixation** | 4% EM grade PFA/PEM, 15min RT | 4% EM grade PFA/PEM, 15min RT |
| **Quenching** | 0.1% sodium borohydride/PBS, 7min RT | 0.1% sodium borohydride/PBS, 7min RT |
| **Blocking** | 10%GS/0.2% Triton/PBS, 1h RT | 10% GS/3% BSA/0.2% Triton/PBS, 1h RT |
| **Primary antibody dilution** | 1:50 in 5% GS/0.1% Triton/PBS, ON at 4°C | 1:50 in blocking serum, ON at 4°C |
| **Washing steps** | All in 0.01M PBS | All in 0.01M PBS |
| **Secondary antibody** | Goat anti-mouse<br>AF647 Plus (A32728) or AF647 (A21236) | Goat anti-mouse<br>AF647 Plus (A32728) |
| **Secondary antibody dilution** | 1:500 in 5% GS/0.1% Triton/PBS, 1h RT | 1:500 in 3% BSA/PBS, 1h RT |

*all secondary antibodies were purchased from Thermo Fisher. Product numbers are indicated in brackets.

**Supplementary Table S3. Summary table of antibodies used in the manuscript**

| Figure | Primary and secondary antibodies used |
|---|---|
| Figure 1 | a: mouse anti-NfL; goat anti-mouse AF647 Plus<br>b: mouse anti-NfL; goat anti-mouse AF488 Plus<br>c: mouse anti-panNa$_v$; goat anti-mouse AF647 Plus<br>d: mouse anti-ankG; goat anti-mouse AF647 Plus |
| Figure 2 | a,b,c,: rabbit anti-tRFP; goat anti-rabbit AF647<br>d: rabbit anti-tRFP; goat anti-rabbit AF647 or AF488 |
| Figure 3 | a: AF647 anti-Tubulin β3<br>b: AF488 anti-Tubulin β3 |
| Figure 4 | a, b: mouse anti-NfL; goat anti-mouse AF647 Plus |
| Figure 5 | a, b: mouse anti-ankG; goat anti-mouse AF647 Plus<br>c, d: mouse anti-panNa$_v$; goat anti-mouse AF647 Plus |
| Figure 6 | a: mouse anti-panNa$_v$; goat anti-mouse AF647<br>b: mouse anti-panNa$_v$; goat anti-mouse AF647 Plus |
| Figure S1 | a: AF647 anti-Tubulin β3<br>b: mouse anti-TUBB3; goat anti-mouse AF647 Plus |
| Figure S2 | a: mouse anti-NfL; goat anti-mouse AF647 Plus<br>b: mouse anti-NfL; goat anti-mouse AF647<br>c: mouse anti-panNa$_v$; goat anti-mouse AF647 Plus<br>d: mouse anti-panNa$_v$; goat anti-mouse AF647 |
| Figure S3 | a: mouse anti-NfL; goat anti-mouse AF647 Plus<br>b: mouse anti-NfL; goat anti-mouse AF647<br>c: mouse anti-NfL; goat anti-mouse AF488 Plus |
| Figure S4 | a: mouse anti-NfL; goat anti-mouse AF555<br>b: mouse anti-NfL; goat anti-mouse AF633 |
| Figure S5 | a, b: rabbit anti-tRFP; goat anti-rabbit AF647<br>c, d: rabbit anti-tRFP; goat anti-rabbit AF488 |
| Figure S6 | a,b: Phalloidin-AF647 |
| Figure S7 | a: mouse anti-NfL; goat anti-mouse AF647 Plus<br>b: mouse anti-NfL; goat anti-mouse AF647 |

### Supplementary Material and Methods

#### Cell culture

For experiments, ND7/23 cells were seeded on a four-well Lab-Tek II chambered #1.5 German coverglass (Thermo Fisher Scientific, cat. no. 155382), 70,000 cells per well. Lab-Teks were coated with poly-D-lysine (Sigma Aldrich, cat. no. P6407) diluted in ddH2O at a 10 µg/ml concentration, and incubated at least 4 h at room temperature (RT). Before seeding wells were washed twice with Dulbecco's phosphate-buffered saline (DPBS; Thermo Fisher Scientific, cat. no. 14190169) or ddH2O and left in the hood to dry completely. Cells were passaged every 2-3 days up to passage 18-20 and were not differentiated.

#### Imaging of actin (phalloidin AF647), NfL and tubulin labelled cells

Phalloidin AF647, neurofilament light chain (NfL) and tubulin labelled cells were first checked in PBS, using fluorescent light source (Lumencor Sola SE II) and Cy5 filter cube or 488 filter cube with 10 ms exposure time and 5 % excitation light intensity. Up to 40 fields of view (positions) per well were picked and xyz coordinates of each field of view were saved in NIS-Elements AR software. Widefield image acquisition was performed automatically by using NIS-Elements ND multipoint acquisition module at 30 ms exposure time and 10 % of fluorescent light source. Both, brightfield and fluorescence images were acquired. Afterwards, PBS was replaced with Vectashield, left 2 minutes to completely cover the sample and acquisition was repeated. Upon the addition of Vectashield we noticed a change in focus, so before acquiring images in Vectashield, every field of view needed to be manually refocused. During the imaging, autoscaling look-up table (LUT) was on, which allowed us to see what would approximately be the correct focus plane, despite the quenching of fluorescent signal.

#### Imaging media for dSTORM

25% Vectashield was made by mixing Vectashield (VS) with Tris-Glycerol in 1:4 v/v ratio. Tris-glycerol was obtained by adding 5 % v/v 1 M Tris, pH 8 to glycerol (Sigma Aldrich, cat. no. G2025), as described previously[1]. For preparation of GLOX buffer, we used Buffer A (10 mM Tris, pH 8, 50 mM NaCl), Buffer B (50 mM Tris, pH 8, 10 mM NaCl) and GLOX solution. GLOX solution was made by mixing 14 mg glucose oxidase (Sigma Aldrich, cat. no. G2133), 50 µl catalase (17 mg/ml; Sigma Aldrich, cat. no. C3155) and 200 µl Buffer A. Aliquots of GLOX solution were kept at -20 °C. GLOX buffer was prepared fresh, prior to use by mixing 7 µl GLOX solution with 690 µl Buffer B containing 10 % w/v glucose (Sigma Aldrich, cat. no. D9559) on ice. For 3D STORM imaging 7 ul of β-mercaptoethanol (Sigma Aldrich, cat. no. M3148) were added to GLOX buffer (GLOX BME).

#### Image analysis and intensity measurements

For the purpose of analysis, we made a macro that allowed us to stack and align all images of each field of view and perform the intensity measurements in same regions of interest (ROIs). Images from mCherry channel were used for alignment and tresholding, while intensity measurements were done in images from AF647 and AF488 channels, respectively.

In all of our experiments (except recovery of AF647) each field of view was imaged two times: in 561 (mCherry) and either 647 (AF647) or 488 (AF488) channel, before and after medium change. As result, for each field of view we had four fluorescent pictures, mCherry and AF647(or AF488) before and after medium change. In AF647 fluorescence recovery experiments each field of view was imaged three times: in mCherry and AF647 channel, before medium change, after changing to Vectashield and after washing in PBS. Consequently, for each field of view in recovery experiments we had six fluorescent pictures, mCherry and AF647 before medium change, in Vectashield and after washing in PBS.

For the purpose of image analysis and intensity measurements we designed macro to open and stack images from mCherry channel, align them using MultiStackReg (developed by Brad Busse http://bradbusse.net/sciencedownloads.html) and TurboReg plugin[2], and save the alignment information in a form of transformation matrices. After that, macro was opening and stacking images from AF647 channel. AF647 images were aligned using transformation matrices that were saved after alignment of mCherry images. This way we ensured that images from 647 channel would be properly aligned even in the case when one of them has very low intensity (e.g. images taken in Vectashield). After alignment both mCherry and AF647 image stacks were cropped to exclude empty space. Same procedure was done for AF488 images.

Region of interest for intensity measurements was created by thresholding in mCherry image stack, using Otsu Dark thresholding algorithm. Thresholding was performed in the image that was taken after medium change to exclude cells that were washed away during the change of imaging media. Resulting ROI contained fluorescently labelled nuclei and we refer to it as nuclear ROI. At this point, user intervention was required, to choose ROI that contains no cells, only background noise (background ROI). Afterwards, intensity measurements (mean intensity and integrated density) of nuclear and background ROIs were performed in AF647 and AF488 stacks, respectively.

As a result of analysis, macro saved intensity measurement results (in a form of a text file), mCherry before/after image stacks, AF488 or AF647 before/after image stacks and nuclear ROIs. Oversaturated images, poorly aligned images and images where all cells were washed away during imaging media change were excluded from further analysis.

Text files with intensity measurement results were imported in Excel and corrected total cell fluorescence (CTCF) was calculated.

CTCF = Integrated Density of nuclear ROI – (area of nuclear ROI x mean fluorescence of the background)

In addition to CTCF, we calculated average fluorescence by dividing CTCF with the area of nuclear ROI. Calculated values were exported to a separate Excel file for statistical analysis.

### Supplementary Details on Statistical Analysis

**Preparation of data analysis based on pilot work**

The dependent variable was the difference (delta) between the fluorescence intensities before and after media change, where the starting medium is always PBS and the change is represented by removing PBS and adding another medium (delta intensity = (intensity PBS – intensity new medium)). Based on prior assumptions, this difference will become minimal in the control condition, where PBS is first removed and the added new medium is again PBS. If another medium is added, delta will be larger and take on an either positive or negative value, depending on whether the intensity in the new medium is smaller or larger than the intensity of PBS.
The dependent variable (delta intensity) delivers continuous values and is on interval scale level, which is a prerequisite to carry out an analysis of variance (ANOVA). The independent variables (factors) were different dyes (AF647 and AF488), different imaging media (PBS, Vectashield, 25 % Vectashield, GLOX) and different experiments (1, 2 and 3).

**Power calculations, randomization and design**

Prior to the actual experiments, it was necessary to carry out power calculations to specify the optimal sample size. These analyses needed to take into account multiple analyses, including the different experiments and post-hoc comparisons. Our main focus was to compare delta intensities between different imaging media applying ANOVA. Our defined type 1 error level was 0.05 (including two sided testing for post-hoc, planned comparisons). We aimed for a power of 0.95 (which equals a type 2 error level of 0.05). Based on pilot work, we had estimated to find at least an effect size between 0.45 and 0.5 and more realistic an effect size between 0.8 and 0.9. For the lowest effect of 0.45 we would need 23 measurements per condition, for the large effect size of 0.9 we would only need 7 measurements per group to reach the pre-specified power of 0.95. To fulfil the assumptions of ANOVA and post-hoc tests in terms of required distributions, we decided for 20 measurements per condition, which lies within the range of 7 to 23 measurements. We decided in favour of a value at the upper limit of this range because the smaller the sample size, the more likely are violations to the assumptions of parametric tests such as ANOVA. We did not go beyond this range, though, to avoid having over-powered statistical tests, where tiniest differences would become statistically significant.
In our pilot work, we also realized that if images are taken truly at random, a minority of the images could not be measured because no cells appeared after changing media (they got washed away), or some of the images were oversaturated. Because randomization was an absolute requirement for our statistical tests, we needed to have a bigger number than 20 images randomly taken to get a total of 20 measurements. To be safe, we decided a priori that 30 images are taken at random and the first 20 valid images (i.e. those that contained cells before and after media change and that did not contain oversaturated pixels) were included in the analysis. The remaining images were not considered in the analysis. This resulted in the following design:
In each experiment, 20 delta intensities were measured for all 4 different imaging media (20 * 4 = 80). Because they were measured for both dyes, AF647 and AF488, there were 160 delta intensities (80 * 2) per experiment. Because we carried out 3 experiments, there were 480 delta intensities in total (160 * 3).
As a result, there was a 2 (dye) * 4 (imaging media) * 3 (experiment) ANOVA on delta intensities. Following our a priori assumptions, we assumed the two dyes would deliver different delta

intensities. In addition, we assumed the 4 imaging media would differ in terms of their delta intensities. Because each experiment took place on a different day with new cells (that were freshly transfected and labelled, etc.), we also expected that the 3 experiments would differ in terms of their delta intensities. In case the 3 experiments indeed differ, different ANOVAs are necessary for each experiment separately, in order to show that the differences between the imaging media point in the same direction and the same post-hoc comparisons are significant in each individual experiment. In these individual ANOVAs, the delta intensities for all imaging media are directly compared to each other for each experiment separately and for each dye separately. Planned post-hoc comparisons (Bonferroni) are subsequently carried out to demonstrate that all post-hoc analyses show the same type of significant differences between the compared imaging media. These planned comparisons correct the significance levels for multiple tests, i.e. they avoid an accumulation of statistical error which would otherwise occur if multiple significance tests were carried out. To give us maximum trust in our results, we chose the strictest of all post-hoc comparisons, which is the Bonferroni correction. Bonferroni delivers the most conservative results, i.e. the least likely to become statistically significant.

**Deriving hypotheses based on prior observations**

Based on pilot experiments, our primary focus was dye AF647, because only in AF647 we noticed an intensity drop in image medium Vectashield compared to PBS. We expected that this drop was shown in all 3 experiments. We did not expect such a drop for dye AF488. We do, however, also expect intensity changes on other imaging media for dye AF488. Based on pilot work, we expected medium sized effects relating to a delta intensity increase for Vectashield and 25 % Vectashield. If this increase is robust, it should appear in all three experiments. This also requires planned post-hoc comparisons (Bonferroni), to demonstrate that the significant differences between the compared imaging media are the same for all 3 experiments.

Based on pilot data, we had yet another hypothesis. We realized an intensity drop seen for dye AF647 after adding Vectashield and found out that the intensity will recover after washing. This would be in contrast to the previously assumed dye cleavage effect, where no recovery would be expected. In order to test whether the AF647 intensity drop seen in Vectashield can actually recover, we carried out a repeated measurement ANOVA. In this ANOVA, we compared 3 intensities (1. at PBS baseline, 2. after removing PBS and adding Vectashield, 3. after replacing Vectashield with PBS and waiting 2.5 hours in PBS imaging medium).

**Data analyses**

In a first step, we carried out a 2 (dye) * 4 (imaging media) * 3 (experiment) ANOVA with the dependent variable delta intensities. There was a significant main effect for dye, $F(1, 456)=819.14$, $p<0.001$, effect size partial $\eta^2=0.64$, Mean AF647=3596.45, Mean AF488=-2569.74, SE(Means)=152.34. Similarly, there was a significant main effect for imaging media, $F(3, 456)=424.15$, $p<0.001$, effect size partial $\eta^2=0.74$, Mean PBS=420.74, Mean Vectashield=6239.29, Mean_25% Vectashield=-4583.83, Mean GLOX=-22.79, SE(Means)=215.45. In addition, there was a significant main effect for experiment, $F(2, 456)=17.58$, $p<0.001$, effect size partial $\eta^2=0.07$, Mean expt 1=864.28; Mean expt 2=-383.07; Mean expt 3=1058.85, SE(Means)=186.58. Finally, all possible combinations of interactions between the factors were significant, $p<0.001$. In summary, the delta intensity values differ between dyes, between imaging media, between experiments and the combination of these factors influence each other differently depending on the chosen combination. Consequently, this analysis including all possible factors and their combinations does not permit conclusions, calling for separate ANOVAs in each of the two dyes and separate ANOVAs in each of the 3 experiments. From a biological point of view, it would also not make sense to carry out one ANOVA across both dyes and all conditions.

**Separate ANOVA for AF647 in each of the 3 experiments**

For dye AF647, there was a main effect for imaging media in all 3 experiments, Experiment 1: $F(3, 76)=164.46$, $p<0.001$, effect size partial $\eta^2=0.87$; Experiment 2: $F(3, 76)=182.45$, $p<0.001$, effect size partial $\eta^2=0.88$; Experiment 3: $F(3, 76)=114.06$, $p<0.001$, effect size partial $\eta^2=82$. Looking at the planned, post-hoc comparisons (Bonferroni), the same result was shown in all three experiments: there was a significant ($p<0.05$) drop of delta intensity from PBS to Vectashield (PBS$\rightarrow$VS): Experiment 1 mean delta intensity drop=19348.86, SE(mean intensity drop)=1150.76, Experiment 2 mean delta intensity drop=10163.16, SE(mean intensity drop)=564.35, Experiment 3 mean delta intensity drop=13821.93, SE(mean intensity drop=975.89). The other media (25 % Vectashield and GLOX) did not differ significantly from PBS in all 3 experiments, i.e. there was no significant intensity drop. Consequently, the results in all 3 experiments support the notion of an intensity drop only from PBS to Vectashield. This was the result we had expected based on our pilot data. The results are robust because we did not rely on experiment 1 alone, but replicated the results in experiments 2 and 3. In all 3 experiments, exactly the same main effects were observed and exactly the same post-hoc comparisons reached statistical significance.

**Analysis of AF647 recovery**

In addition, we carried out a repeated measures ANOVA to test whether the intensity drop seen in Vectashield can actually recover. We had hypothesized this recovery based on earlier observations we had made in our laboratory when running pilot experiments. Now, we could not rely on delta intensities from 2 measurements, but were rather interested in intensities at 3 different time points. We compared three intensities (1. at PBS baseline, 2. after removing PBS and adding Vectashield, 3. after replacing Vectashield with PBS and waiting for 2.5 hours in PBS imaging medium). We again performed these comparisons for all three experiments separately. In this example, we had a repeated measurement with three consecutive measurements. In this case, the respective analysis depends on a prerequisite for a repeated measurements ANOVA: if sphericity can be assumed based on the Mauchly's Test of Sphericity, no corrections need to be performed. Otherwise, the Greenhouse-Geisser Test of an overall "within subjects effect" with corrected degrees of freedom has to be applied. Because the Mauchly's Test of Sphericity revealed a significant result for all three experiments ($p<0.001$), sphericity could not be assumed and the Greenhouse-Geisser Test was applied for all three experiments. In all three experiments, intensity differences between the 3 repeated conditions were statistically significant (1. PBS baseline, 2. after removing PBS and adding Vectashield, 3. after replacing Vectashield with PBS and waiting for 2.5 hours in PBS imaging medium), Experiment 1: $F(1.01, 19.26)=162.68$, $p<0.001$; effect size partial $\eta^2=0.895$; Experiment 2: $F(1.03, 19.48)=175.17$, $p<0.001$, effect size partial $\eta^2=0.9$; Experiment 3: $F(1.18, 22.38)=104.28$, $p<0.001$, effect size partial $\eta^2=0.85$. Given that all three experiments showed a significant overall effect for the repeated conditions, we carried out planned, post-hoc tests with Bonferroni correction to evaluate which of the three time points significantly differed from each other. In each of the three experiments, there was a significant intensity drop from 1. PBS baseline to 2. after removing PBS and adding Vectashield (PBS→VS; in each experiment, $p<0.05$). Similarly, there was a significant recovery from 2. after removing PBS and adding Vectashield to 3. After replacing Vectashield with PBS and waiting for 2.5 hours in PBS imaging medium (VS→PBS:recovery, in each experiment, $p<0.05$). In all three experiments, the recovery was never as strong as to reach the original intensity (in each experiment, $p<0.05$). Consequently, the results demonstrating the recovery effect were very robust. The drop as well as the recovery effect was not only shown in Experiment 1, but replicated exactly in Experiments 2 and 3. In addition, all other post-hoc tests revealed the same results in all three experiments.

Detailed results are as follows:

Experiment 1 had a PBS baseline intensity mean=24036.37, SE(mean)=1928.87, a drop after removing PBS and adding Vectashield with a mean=3638.06, SE(mean)=359.85, and a recovery after replacing Vectashield with PBS and waiting for 2.5 hours with a mean=16554.7, SE(mean)=1394.89.

Experiment 2 had a PBS baseline intensity mean=12270.01, SE(mean)=921.22, a drop after removing PBS and adding Vectashield with a mean=1855.48, SE(mean)=151.16, and a recovery after replacing Vectashield with PBS and waiting for 2.5 hours with a mean=9834.3, SE(mean)=758.21.

Experiment 3 had a PBS baseline intensity mean=16984.29, SE(mean)=1589.18, a drop after removing PBS and adding Vectashield with a mean=2376.21, SE(mean)=247.32, and a recovery after replacing Vectashield with PBS and waiting for 2.5 hours with a mean=13369.76, SE(mean)=1367.025.

**Separate ANOVA for AF488 in each of the 3 experiments**

For dye AF488, there was also a main effect for imaging media in all 3 experiments, Experiment 1: $F(3, 76)=144.36$, $p<0.001$, partial $\eta^2=0.85$; Experiment 2: $F(3, 76)=160.66$, $p<0.001$, partial $\eta^2=0.86$; Experiment 3: $F(3, 76)=101.86$, $p<0.001$, partial $\eta^2=0.8$. Looking at the planned, post-hoc comparisons (Bonferroni), the same result was demonstrated in all three experiments: both Vectashield (PBS→VS) and 25 % Vectashield (PBS→25% VS) showed an increase in delta intensity, which was significantly different from the delta intensity of PBS and GLOX ($p<0.05$), while the delta intensities of PBS and GLOX did not differ significantly from each other. From our pilot work, we had assumed a medium effect size. We did not expect an effect as big as it turned out in our analysis, which came as a surprise to us. The effect was already big in experiment 1 and replicated in experiments 2 and 3. In detail, the results were as follows:

In experiment 1, there were no significant delta intensity differences between both PBS imaging media and between PBS and GLOX, but delta intensity showed an increase from PBS to Vectashield with a mean=3906.7 and an increase from PBS to 25 % Vectashield with a mean=9579.34, SE(means)=400.53 (with post-hoc Bonferroni $p<0.05$). The increase was stronger for 25% Vectashield than for Vectashield (with post-hoc Bonferroni $p<0.05$)

In experiment 2, there were no significant delta intensity differences between both PBS imaging media and between PBS and GLOX, and delta intensity showed an increase from PBS to Vectashield with a mean=1781.265 and an increase from PBS to 25 % Vectashield with a mean=10930.1, SE(means)=415.99 (with post-hoc Bonferroni $p<0.05$). The increase was once again stronger for 25 % Vectashield than for Vectashield (with post-hoc Bonferroni $p<0.05$).

In experiment 3, there were once again no significant delta intensity differences between both PBS imaging media and between PBS and GLOX, and delta intensity showed an increase from PBS to Vectashield with a mean=2297.175 and an increase from PBS to 25 % Vectashield with a mean=3959.1, SE(means)=199.99 (with post-hoc Bonferroni $p<0.05$). The increase was again stronger for 25 % Vectashield than for Vectashield (with post-hoc Bonferroni $p<0.05$).
